# SARS-CoV-2 remodels the Golgi apparatus to facilitate viral assembly and secretion

**DOI:** 10.1101/2022.03.04.483074

**Authors:** Jianchao Zhang, Andrew Kennedy, Daniel Macedo de Melo Jorge, Lijuan Xing, Whitney Reid, Sarah Bui, Joseph Joppich, Molly Rose, Sevval Ercan, Qiyi Tang, Andrew W. Tai, Yanzhuang Wang

## Abstract

The COVID-19 pandemic is caused by SARS-CoV-2, an enveloped RNA virus. Despite extensive investigation, the molecular mechanisms for its assembly and secretion remain largely elusive. Here, we show that SARS-CoV-2 infection induces global alterations of the host endomembrane system, including dramatic Golgi fragmentation. SARS-CoV-2 virions are enriched in the fragmented Golgi. Disrupting Golgi function with small molecules strongly inhibits viral infection. Significantly, SARS-CoV-2 infection down-regulates GRASP55 but up-regulates TGN46 protein levels. Surprisingly, GRASP55 expression reduces both viral secretion and spike number on each virion, while GRASP55 depletion displays opposite effects. In contrast, TGN46 depletion only inhibits viral secretion without affecting spike incorporation into virions. TGN46 depletion and GRASP55 expression additively inhibit viral secretion, indicating that they act at different stages. Taken together, we show that SARS-CoV-2 alters Golgi structure and function to control viral assembly and secretion, highlighting the Golgi as a potential therapeutic target for blocking SARS-CoV-2 infection.

## Introduction

Since December 2019, an unprecedented pandemic caused by coronavirus disease 2019 (COVID-19) has raised a serious threat to human health worldwide. SARS-CoV-2 (severe acute respiratory syndrome coronavirus 2), the etiologic agent for COVID-19, is an enveloped, single-stranded positive-sense RNA virus in the beta-coronavirus family. Vaccination is the most critical and effective strategy to prevent COVID-19 from spreading. However, vaccines are not effective in all individuals, such as the immunocompromised, and the possible emergence of SARS-CoV-2 variants with immune evasion and/or increased pathogenicity continues to pose a great threat to the existing vaccines (Planas et al., 2021). Therefore, it is urgent and important to elucidate the detailed mechanisms of the SARS-CoV-2 infection cycle and thus develop various strategies to block the replication and spread of all variants.

Entry into host cells is the first step of virus infection and acts as a determinant of viral infectivity and pathogenesis. Since the pandemic outbreak, the cell entry mechanisms of SARS-CoV-2 have been extensively explored. Like SARS-CoV-1, SARS-CoV-2 employs human angiotensin-converting enzyme 2 (ACE2) as the entry receptor. The spike (S) protein on the virus interacts with ACE2 at the host cell surface, allowing the virus to attach to the host cell membrane. After cleavage by the transmembrane serine protease 2 (TMPRSS2), the spike protein is activated and initiates a fusion process of the viral membrane with the host cell plasma membrane. When TMPRSS2 is absent, SARS-CoV-2 is endocytosed and the spike protein is activated by Cathepsin B and L (CatB/L) inside the lysosomal lumen (Hoffmann et al., 2020a). Compared to SARS-CoV-1, the spike protein of SARS-CoV-2 possesses a unique furin cleavage site that facilitates its cell entry and contributes to its higher infectivity (Shang et al., 2020). Following entry, the genomic RNA is released into the cytoplasm. The polyproteins pp1a and pp1ab, translated from two large open reading frames ORF1a and ORF1b, are self-processed into 16 individual non-structural proteins (Nsps), which modulate the endoplasmic reticulum (ER) to form double-membrane vesicles (DMVs) where the viral replication and transcription complex (RTC) is anchored to produce abundant viral RNAs (Khoury et al., 2020).

In addition to the 16 Nsps, the SARS-CoV-2 genome also encodes four structural (S, M, E, and N) and 9 accessory proteins (Gordon et al., 2020). Three of the four viral structural proteins (S, E, and M) are synthesized in the ER and transported to the ER-to-Golgi intermediate compartment (ERGIC), where virions are assembled and budded into the lumen of the ERGIC and Golgi. In the ER and Golgi, the spike protein is highly glycosylated (Sanda et al., 2021; Tian et al., 2021), which affects its folding, stability, interaction with ACE2, and susceptibility to vaccines. Integrative imaging revealed that SARS-CoV-2 infection triggers Golgi fragmentation (Cortese et al., 2020), but the cause and consequence of Golgi fragmentation remain largely elusive.

It was generally believed that mature viral particles of several coronaviruses, including SARS-CoV-1, undergo conventional protein secretory pathways for release, (Siu et al., 2008; Westerbeck and Machamer, 2015, 2019), which is also true for SARS-CoV-2 (Eymieux et al., 2021). However, some other reports indicated that SARS-CoV-2 virus exits host cells in a Golgi-independent manner by lysosomal exocytosis (Chen et al., 2021; Ghosh et al., 2020). Thus, whether SARS-CoV-2 traffics through the Golgi for release into the extracellular space is still controversial.

Here, we systematically investigated the interaction between SARS-CoV-2 and the host endomembrane system. Our study revealed that SARS-CoV-2 infection causes a dramatic disruption of the Golgi structure, and disruption of Golgi homeostasis by small molecules significantly reduces viral infection. By surveying a large number of Golgi proteins, we found that SARS-CoV-2 infection reduces GRASP55 but increases TGN46 protein level. Importantly, expression of GRASP55 or depletion of TGN46 in cells significantly inhibits viral infection and secretion. Interestingly, manipulation of GRASP55 also affects spike number on each virion. Therefore, our study uncovered a novel mechanism by which SARS-CoV-2 remodels the Golgi structure and function to facilitate viral assembly and secretion, suggesting the Golgi as a potential drug target for COVID-19 treatment.

## Results

### SARS-CoV-2 infection induces dramatic morphological alterations of multiple subcellular organelles, in particular the Golgi

Enveloped viruses often utilize the host secretory pathway to assemble and bud out of the cell (Welsch et al., 2007). How SARS-CoV-2, an enveloped beta-coronavirus, modulates the endomembrane system for its assembly and exit is an important but unanswered question. To address this question, we performed morphological analyses of multiple subcellular organelles in the secretory pathway, including the ER, ERGIC, Golgi, autophagosome, endosome, and lysosome, in SARS-CoV-2-infected Huh7-ACE2 cells, a permanent hepatoma cell line that is often used in studies of SARS-CoV-2 infection (Sherman et al., 2021). After 24 h infection by SARS-CoV-2 (USA-WA1 strain, same as below unless specified), the ER structure marked by calreticulin was not dramatically altered (Table 1, Figure S1A-C). ERGIC53 was normally concentrated in the perinuclear region in uninfected cells, while in SARS-CoV-2 infected cells it displayed as dispersed puncta distributed in the cytoplasm (Figures 1A-B). Further quantitation of the images indicated that the total area of ERGIC53 did not change (Figure 1C), and its relative expression level was modestly reduced (Figure 1D).

**Figure 1.**
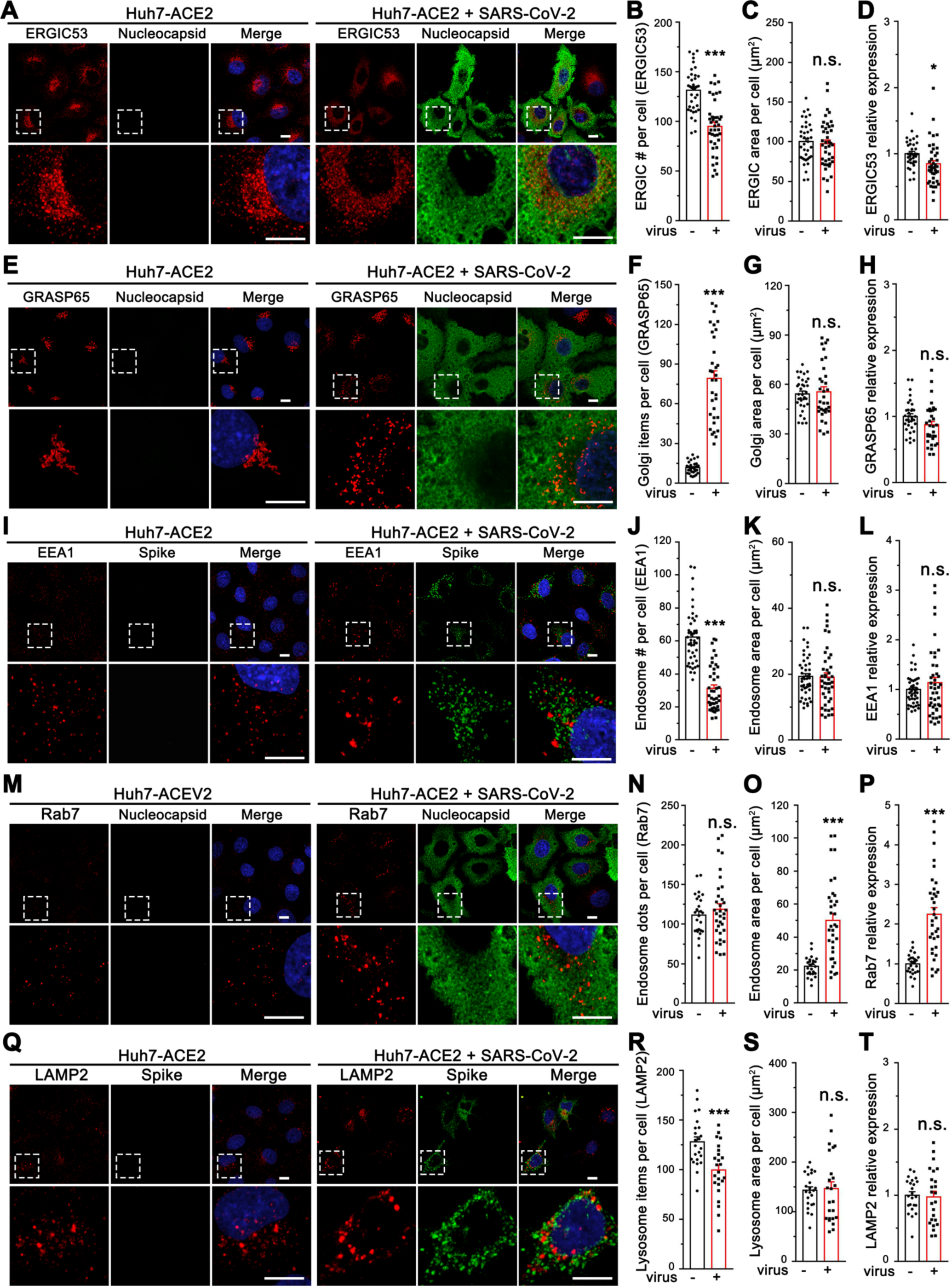
SARS-CoV-2 infection induces global morphological changes of host cell membrane organelles. (A) Representative confocal images of Huh7-ACE2 cells incubated with or without SARS-CoV-2 (MOI = 1) for 24 h and stained for ERGIC53 and nucleocapsid. (B-D) Quantification of ERGIC53 for the number of puncta (B), area (C), and relative expression (D) in A. (E) Representative confocal images of Huh7-ACE2 cells incubated with or without SARS-CoV-2 (MOI = 1) for 24 h and stained for a cis-Golgi marker GRASP65 and nucleocapsid. (F-H) Quantification of GRASP65 item number (F), area (G), and relative expression (H) in E. (I) Representative confocal images of Huh7-ACE2 cells incubated with or without SARS-CoV-2 (MOI = 1) for 24 h and stained for an early endosome marker EEA1 and spike. (J-L) Quantification of EEA1 item number (J), area (K), and relative expression (L) in I. (M) Representative confocal images of Huh7-ACE2 cells incubated with or without SARS-CoV-2 (MOI = 1) for 24 h and stained for a late endosome marker Rab7 and nucleocapsid. (N-P) Quantification of Rab7 in M. (Q) Representative confocal images of Huh7-ACE2 cells incubated with or without SARS-CoV-2 (MOI = 1) for 24 h and stained for a late endosome/lysosome marker LAMP2 and spike. (R-T) Quantification of Rab7 in Q. Boxed areas in the upper panels are enlarged and shown underneath. Scale bars in all panels, 10 μm. All quantitation data are shown as mean ± SEM from three independent experiments. Statistical analyses are performed using two-tailed Student’s t-test. *, p < 0.05; **, p < 0.01; ***, p < 0.001; n.s., not significant.

**Table 1.**
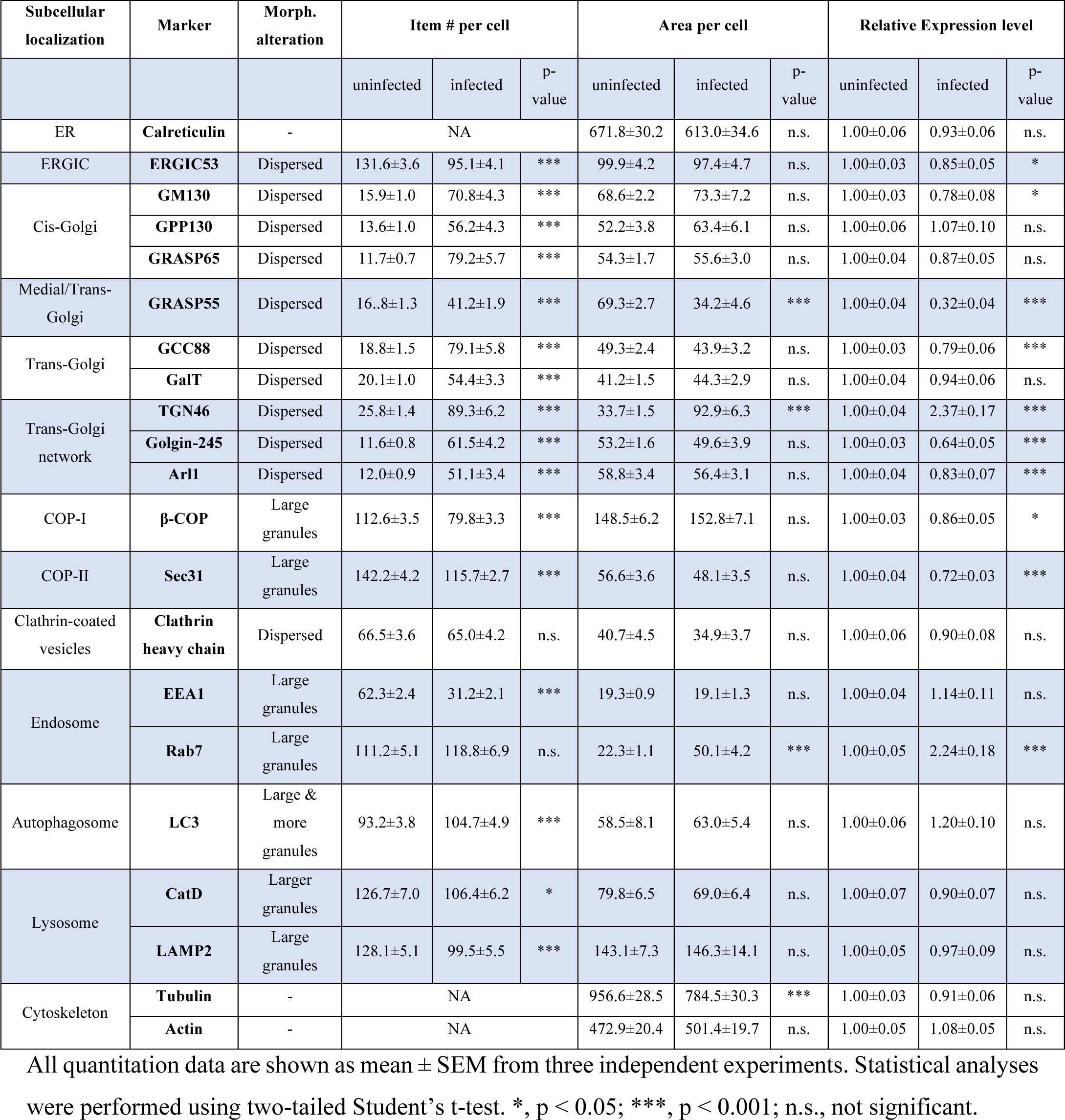
Summary of morphological changes of subcellular organelles after SARS-CoV-2 infection. All quantitation data are shown as mean ± SEM from three independent experiments. Statistical analyses were performed using two-tailed Student’s t-test. *, p < 0.05; ***, p < 0.001; n.s., not significant.

| Subcellular localization | Marker | Morph. alteration | Item # per cell |  |  | Area per cell |  |  | Relative Expression level |  |  |
| --- | --- | --- | --- | --- | --- | --- | --- | --- | --- | --- | --- |
|  |  |  | uninfected | infected | p-value | uninfected | infected | p-value | uninfected | infected | p-value |
| ER | **Calreticulin** | - | NA |  |  | 671.8±30.2 | 613.0±34.6 | n.s. | 1.00±0.06 | 0.93±0.06 | n.s. |
| ERGIC | **ERGIC53** | Dispersed | 131.6±3.6 | 95.1±4.1 | *** | 99.9±4.2 | 97.4±4.7 | n.s. | 1.00±0.03 | 0.85±0.05 | * |
| Cis-Golgi | **GM130** | Dispersed | 15.9±1.0 | 70.8±4.3 | *** | 68.6±2.2 | 73.3±7.2 | n.s. | 1.00±0.03 | 0.78±0.08 | * |
|  | **GPP130** | Dispersed | 13.6±1.0 | 56.2±4.3 | *** | 52.2±3.8 | 63.4±6.1 | n.s. | 1.00±0.06 | 1.07±0.10 | n.s. |
|  | **GRASP65** | Dispersed | 11.7±0.7 | 79.2±5.7 | *** | 54.3±1.7 | 55.6±3.0 | n.s. | 1.00±0.04 | 0.87±0.05 | n.s. |
| Medial/Trans-Golgi | **GRASP55** | Dispersed | 16.8±1.3 | 41.2±1.9 | *** | 69.3±2.7 | 34.2±4.6 | *** | 1.00±0.04 | 0.32±0.04 | *** |
| Trans-Golgi | **GCC88** | Dispersed | 18.8±1.5 | 79.1±5.8 | *** | 49.3±2.4 | 43.9±3.2 | n.s. | 1.00±0.03 | 0.79±0.06 | *** |
|  | **GalT** | Dispersed | 20.1±1.0 | 54.4±3.3 | *** | 41.2±1.5 | 44.3±2.9 | n.s. | 1.00±0.04 | 0.94±0.06 | n.s. |
| Trans-Golgi network | **TGN46** | Dispersed | 25.8±1.4 | 89.3±6.2 | *** | 33.7±1.5 | 92.9±6.3 | *** | 1.00±0.04 | 2.37±0.17 | *** |
|  | **Golgin-245** | Dispersed | 11.6±0.8 | 61.5±4.2 | *** | 53.2±1.6 | 49.6±3.9 | n.s. | 1.00±0.03 | 0.64±0.05 | *** |
|  | **Arl1** | Dispersed | 12.0±0.9 | 51.1±3.4 | *** | 58.8±3.4 | 56.4±3.1 | n.s. | 1.00±0.04 | 0.83±0.07 | *** |
| COP-I | **β-COP** | Large granules | 112.6±3.5 | 79.8±3.3 | *** | 148.5±6.2 | 152.8±7.1 | n.s. | 1.00±0.03 | 0.86±0.05 | * |
| COP-II | **Sec31** | Large granules | 142.2±4.2 | 115.7±2.7 | *** | 56.6±3.6 | 48.1±3.5 | n.s. | 1.00±0.04 | 0.72±0.03 | *** |
| Clathrin-coated vesicles | **Clathrin heavy chain** | Dispersed | 66.5±3.6 | 65.0±4.2 | n.s. | 40.7±4.5 | 34.9±3.7 | n.s. | 1.00±0.06 | 0.90±0.08 | n.s. |
| Endosome | **EEA1** | Large granules | 62.3±2.4 | 31.2±2.1 | *** | 19.3±0.9 | 19.1±1.3 | n.s. | 1.00±0.04 | 1.14±0.11 | n.s. |
|  | **Rab7** | Large granules | 111.2±5.1 | 118.8±6.9 | n.s. | 22.3±1.1 | 50.1±4.2 | *** | 1.00±0.05 | 2.24±0.18 | *** |
| Autophagosome | **LC3** | Large & more granules | 93.2±3.8 | 104.7±4.9 | *** | 58.5±8.1 | 63.0±5.4 | n.s. | 1.00±0.06 | 1.20±0.10 | n.s. |
| Lysosome | **CatD** | Larger granules | 126.7±7.0 | 106.4±6.2 | * | 79.8±6.5 | 69.0±6.4 | n.s. | 1.00±0.07 | 0.90±0.07 | n.s. |
|  | **LAMP2** | Large granules | 128.1±5.1 | 99.5±5.5 | *** | 143.1±7.3 | 146.3±14.1 | n.s. | 1.00±0.05 | 0.97±0.09 | n.s. |
| Cytoskeleton | **Tubulin** | - | NA |  |  | 956.6±28.5 | 784.5±30.3 | *** | 1.00±0.03 | 0.91±0.06 | n.s. |
|  | **Actin** | - | NA |  |  | 472.9±20.4 | 501.4±19.7 | n.s. | 1.00±0.05 | 1.08±0.05 | n.s. |

The Golgi structure was significantly affected by SARS-CoV-2 infection. In contrast to the well-organized ribbon-like structure in uninfected cells, as labeled by the Golgi protein GRASP65, the Golgi was seen as small fragments dispersed in the entire cytoplasm in infected cells (Figure 1E-F), although the total area and expression level of GRASP65 did not change (Figure 1G-H). In control cells, COPI, COPII, and clathrin coats, indicated by α-COP, Sec31, and clathrin heavy chain, respectively, are accumulated in the perinuclear region, whereas in SARS-CoV-2 infected cells they are also dispersed in the cytoplasm (Figure S1D-O), consistent with Golgi fragmentation.

The endosome/lysosome system was also affected by SARS-CoV-2 infection. The early endosome marker EEA1 and the late endosome marker Rab7 were both found in larger puncta in infected cells (Figures 1I-P), suggesting aggregation of the membranes. While EEA1 expression level did not change (Figures 1L), Rab7 was up-regulated (Figures 1P), suggesting a growth of late endosomes. The aggregation of endosome/lysosome membranes was supported by the late endosome/lysosome marker LAMP2, which displayed fewer but larger puncta in infected cells (Figures 1Q-T). In addition, modest effects were observed for LC3 and Cathepsin D (Figure S2A-H), which represent autophagosome and active lysosomes, respectively. While microtubule cytoskeleton appeared to be partially depolymerized, actin microfilament organization did not seem to change in SARS-CoV-2 infected cells (Figure S2I-N). Taken together, SARS-CoV-2 infection alters the structures of the ERGIC, endosomes, lysosomes, and particularly, the Golgi.

### Disrupting Golgi functions by small molecules inhibits SARS-CoV-2 infection

We reasoned that the morphological changes of the endomembrane system may facilitate SARS-CoV-2 infection, and if so, disrupting the functions of these membrane organelles may impact viral infection. Therefore, we performed a focused small molecule screen to determine whether molecules targeting the endomembrane system could inhibit SARS-CoV-2 infection. Prior to the experiment, we titrated each chemical compound and selected the highest dose that was widely used and did not cause detectable cell death. We then used this concentration to treat cells during SARS-CoV-2 infection. Among a total of 27 drugs tested, 22 drugs displayed significant inhibitory effects on SARS-CoV-2 infection (Figure 2A-B). Three ER stress inducers, thapsigargin (Tg, a SERCA calcium pump inhibitor), tunicamycin (Tm, a protein glycosylation inhibitor), and dithiothreitol (DTT, a reducing agent that disrupts disulfide bonds of proteins), significantly inhibited viral infection. Interestingly, distinct from Tg, ionomycin (Iono), a calcium ionophore that triggers calcium influx across the plasma membrane as well as ER membranes, exhibited no effect on viral infection. Two Golgi function modulators, brefeldin A (BFA, a fungal product that inhibits ER-to-Golgi trafficking) and monensin (an ionophore that inhibits trans-Golgi transport), displayed over 98% inhibition of viral infection, suggesting an important role of the Golgi in SARS-CoV-2 infection.

**Figure 2.**
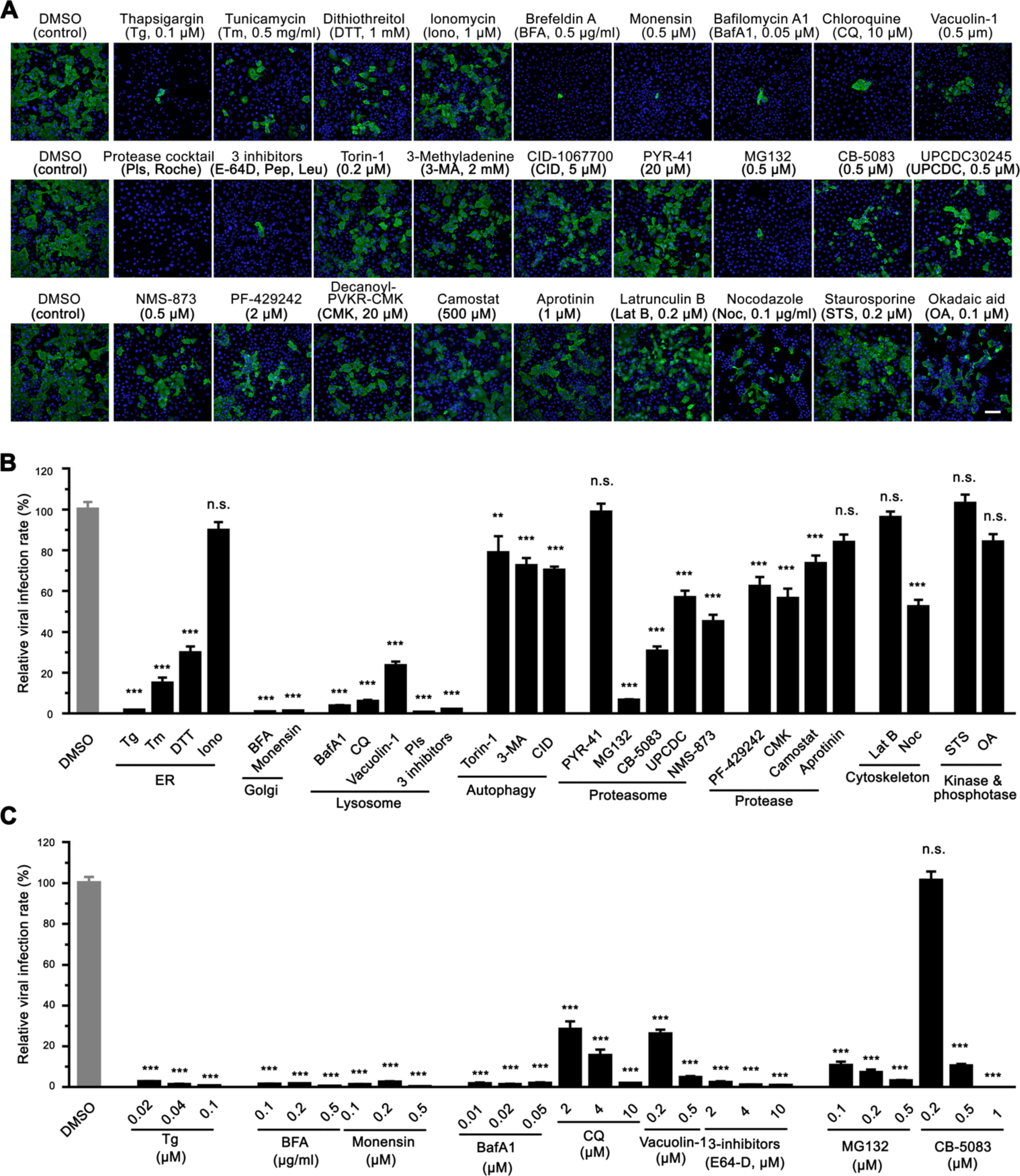
Chemicals disrupting Golgi functions inhibit SARS-CoV-2 infection. (A) Representative immunofluorescence images of Huh7-ACE2 cells infected with SARS-CoV-2 (MOI = 1) for 24 h in the presence of indicated chemicals and stained for nucleocapsid and DNA. The concentrations of the chemical are shown on the figure or as follows: protease inhibitor cocktails (PIs), 100 ml/tablet; 3 inhibitors, 10 μM E-64D, 20 μM pepstatin, 100 μM leupeptin. Scale bar, 100 μm. (B) Quantification of the viral infection rate in A, with the control normalized to 100%. Data are shown as mean ± SEM from 20-30 random images from two or three independent experiments. (C) Quantification of the viral infection rate in the presence of indicated chemicals with different concentrations (images are shown in Figure S3), with the control normalized to 100%. Data are shown as mean ± SEM from 10 representative images. Statistical analyses are performed using one-way ANOVA, Tukey’s multiple comparison test. *, p < 0.05; ***, p < 0.001; n.s., not significant.

### Chemicals interfering with lysosome functions and protein homeostasis inhibit SARS-CoV-2 infection

The lysosome is a key subcellular organelle in both endocytic and exocytic pathways essential for viral entry and secretion. Indeed, disruption of lysosomal function with either bafilomycin A1 (BafA1, an inhibitor of vacuolar-type H^+^-ATPase that blocks autolysosome acidification), chloroquine (CQ, an FDA-approved antimalarial drug that increases the pH of acid vesicles), Vacuolin-1 (a cell-permeable compound that can induce large and swollen lysosomes), a cocktail of protease inhibitors (PIs, which inhibit a broad range of cellular proteases), or three lysosomal protease inhibitors (E64d, leupeptin, and pepstatin, which inhibit lysosomal hydrolases), significantly reduced viral infection (Figure 2A-B).

Interestingly, both autophagy inducer (Torin-1) and autophagy inhibitors, 3-methyladenine (3-MA) and CID 1067700 (CID), displayed similar inhibitory effects on viral infection, although to a lesser degree than lysosomal inhibitors, implying that autophagy may also be involved in SARS-CoV-2 infection as previously suggested (Gorshkov et al., 2021).

The proteostasis network is often hijacked by viruses to satisfy the high demands for viral protein synthesis and folding (Aviner and Frydman, 2020). Therefore, we blocked proteasome-mediated protein degradation and tested the effect on SARS-CoV-2 infection (Figure 2A-B). Inhibition of proteasome-mediated protein degradation with MG132 (a proteasome inhibitor) potently inhibited viral infection, while inhibition of ubiquitin conjugation by PYR-41 (an E1 ubiquitin-activating enzyme inhibitor) displayed no effect. It is possible that a higher concentration of PYR-41 is required to entirely block the first step of ubiquitination; alternatively, the virus may possess the capability to bypass the E1 enzyme like some bacteria (Qiu et al., 2016). Inhibition of the AAA ATPase p97/VCP with CB-5083, UPCDC-30245, or NMS-873 also significantly reduced viral infection, indicating that the proteostasis pathway may serve as a potential target for the treatment of COVID-19. In addition, three protease inhibitors, PF-429242 (a site-1 protease inhibitor), Decanoyl-RVKR-CMK (a furin inhibitor), and camostat mesylate (a TMPRSS2 inhibitor), displayed modest inhibitory effects on SARS-CoV-2 infection, while aprotinin (an inhibitor of serine proteases including trypsin, chymotrypsin, and plasmin) had no effect.

Latrunculin B (LatB, an actin polymerization inhibitor) did not inhibit viral infection, but nocodazole (Noc, a microtubule polymerization inhibitor) exhibited about 50% inhibition of viral infection (Figure 2A-B), consistent with a previous report that the microtubule cytoskeleton is required for SARS-CoV-2 infection (Cortese et al., 2020). In our study, neither staurosporine (STS, a broad-spectrum protein kinase inhibitor) nor okadaic acid (OA, an inhibitor of several serine/threonine phosphatases) showed a significant effect on viral infection (Figure 2A-B).

To confirm these observations, we selected the 9 most effective inhibitors, which target the ER (Tg), Golgi (BFA and monensin), lysosome (BafA1, CQ, Vacuolin-1, and 3-inhibitors), and protein homeostasis (MG132 and CB-5083), respectively, and performed dose-response assays on SARS-CoV-2 infection. All 9 compounds, in particular, Tg, BFA, monensin, BafA1, and 3-inhibitors, dramatically and consistently inhibited SARS-CoV-2 infection even at much lower concentrations (Figure 2C, Figure S3A). To test whether these molecules inhibit viral infection at the entry level, we conducted a SARS-CoV-2 spike pseudotyped virus cell entry assay (Lei et al., 2020). As expected, pseudovirus cell entry depends on ACE2 expression (Figure S3B) and is inhibited by BafA1, CQ, and lysosomal inhibitors, but not BFA. These results highlight the significance of lysosomes, but not the Golgi, in SARS-CoV-2 cell entry (Figure S3C). Taken together, our results suggest that lysosomes play important roles in viral entry, while the Golgi is important for viral infection at other steps.

### SARS-CoV-2 infection alters the Golgi structure

To better understand the role of the Golgi in SARS-CoV-2 infection, we systematically analyzed the Golgi structure in SARS-CoV-2 infected cells by immunofluorescence (IF) and electron microscopy (EM). Using GM130, β-1, 4-Galactosyltransferase 1 (GalT), and Golgin-245 to represent the cis-Golgi, trans-Golgi, and trans-Golgi network, respectively, we found that all sub-compartments of the Golgi were dramatically fragmented after SARS-CoV-2 infection (Figure 3A, F and K). Despite the dispersal of the Golgi in virus-infected cells (Figure 3B-C, 3G-H, and 3L-M), the Golgi area of all three proteins was not altered. While the expression levels of GM130 and GalT remained unchanged or slightly changed, Golgin-245 level was reduced (Figure 3D-E, 3I-J, and 3N-O). Both IF and 3D reconstruction of the Golgi by staining of a cis-Golgi marker GM130 demonstrated that spike was highly enriched in the Golgi fragments in SARS-CoV-2 infected cells (Figure 3A, Movie S1and S2), indicating an important role of the Golgi in viral infection.

**Figure 3.**
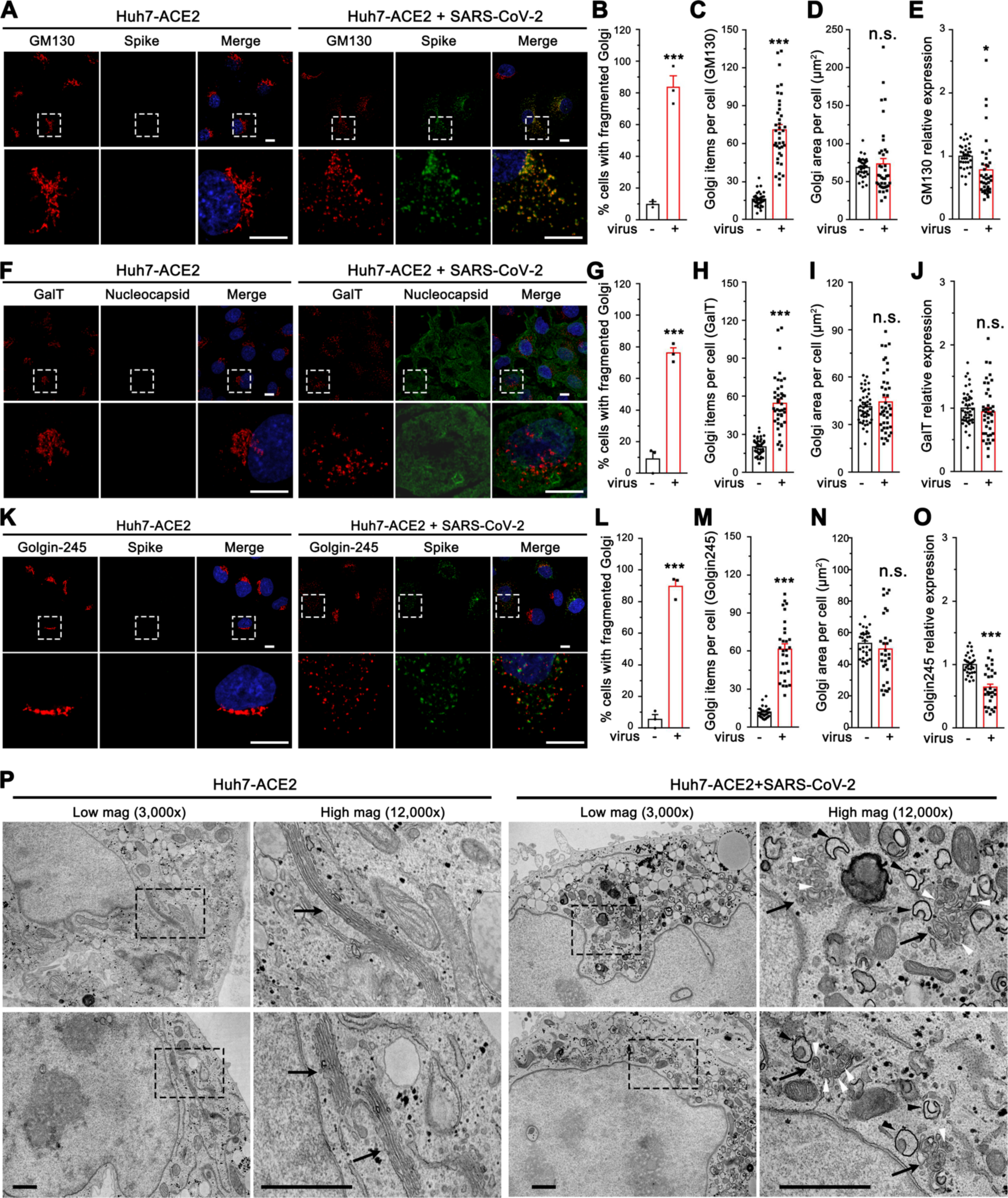
SARS-CoV-2 infection dramatically disrupts the Golgi structure. (A) Representative confocal images of Huh7-ACE2 cells incubated with or without SARS-CoV-2 (MOI = 1) for 24 h and stained for a cis-Golgi marker GM130 and nucleocapsid. (B-E) Quantification of A for the percentage of cells with fragmented Golgi (B), GM130 punctum number (C), area (D), and relative expression level (E). (F) Representative confocal images of Huh7-ACE2 cells incubated with or without SARS-CoV-2 (MOI = 1) for 24 h and stained for a trans-Golgi marker GalT and nucleocapsid. (G-J) Quantification of GalT in F. (K) Representative confocal images of Huh7-ACE2 cells incubated with or without SARS-CoV-2 (MOI = 1) for 24 h and stained for a TGN marker Golgin-245 and spike. Boxed areas in the upper panels of A, F and K are enlarged and shown underneath. Scale bars in all fluorescence images, 10 μm. (L-O) Quantification of Golgin-245 in K. (P) Electron micrographs of Huh7-ACE2 incubated with or without SARS-CoV-2 (MOI = 1) for 24 h under two different magnifications. Boxed areas on the left images are enlarged on the right. Black arrows, white arrowheads, and black arrowheads indicate Golgi membranes, viral particles, and DMVs, respectively. Scale bars, 500 nm. Data are shown as mean ± SEM from three independent experiments. Statistical analyses are performed using two-tailed Student’s t-test. *, p < 0.05; ***, p < 0.001; n.s., not significant.

Under EM, the Golgi displayed dramatically different features between uninfected and SARS-CoV-2 infected cells (Figure 3P, Figure S4). In uninfected cells, the Golgi was highly concentrated in the perinuclear region and displayed long cisterna and stacked structures. In infected cells, the Golgi stacks were severely disorganized and fragmented, with most Golgi membranes vesiculated. Interestingly, many virus particles were observed inside the swollen Golgi lumen, suggesting that SARS-CoV-2 traffics through the Golgi apparatus and causes Golgi fragmentation. Consistent with previous reports (Cortese et al., 2020), we also observed a number of DMVs (Figure 3P, marked by black arrowheads). Taken together, SARS-CoV-2 infection triggers severe Golgi fragmentation.

### SARS-CoV-2 infection down-regulates GRASP55 and up-regulates TGN46 levels

To further investigate the molecular mechanisms of Golgi fragmentation induced by viral infection, we analyzed the morphology and expression level of more Golgi proteins in SARS-CoV-2 infected cells by immunofluorescence microscopy. Consistent with results from the Golgi markers tested above (Figure 1E; Figure 3A, F, and K), GCC88 and Arl1 also showed a remarkably higher Golgi fragmentation frequency and Golgi item number in infected cells (Figure S5A-C and S5F-H), with an unchanged Golgi area and marginally changed expression level (Figure S5D-E and S5I-J). Similarly, the Golgi structure marked by GRASP55 or TGN46 was significantly dispersed with a higher punctum number in infected cells (Figure 4A-C, and 4F-H). However, different from all other Golgi markers tested above, both GRASP55 area and expression level were significantly reduced (Figure 4D-E), whereas both TGN46 area and expression level were dramatically increased after SARS-CoV-2 infection (Figure 4I-J).

**Figure 4.**
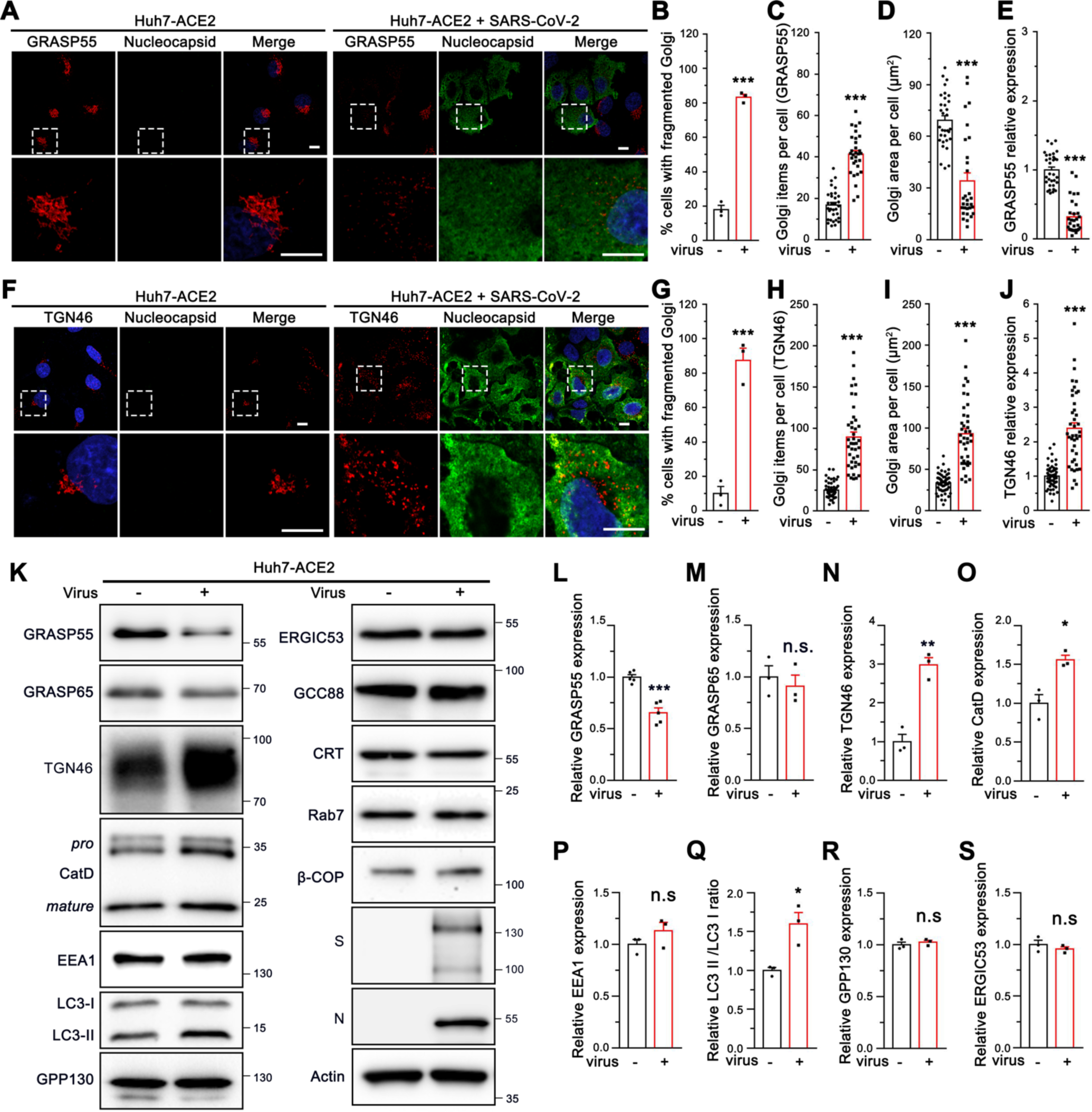
SARS-CoV-2 down-regulates GRASP55 and up-regulates TGN46 expression in Huh7-ACE2 cells. (A) SARS-CoV-2 infection reduces GRASP55 expression. Representative confocal images of Huh7-ACE2 cells incubated with or without SARS-CoV-2 (MOI = 1) for 24 h and stained for GRASP55 and nucleocapsid. (B-E) Quantification of the percentage of cells with fragmented Golgi (B), GRASP55 item number (C), area (D), and relative expression level (E) in A. (F) SARS-CoV-2 infection increases TGN46 expression. Representative confocal images of Huh7-ACE2 cells incubated with or without SARS-CoV-2 (MOI = 1) for 24 h and stained for TGN46 and nucleocapsid. Boxed areas in the upper panels of A and F are enlarged and shown underneath. Scale bars, 10 μm. (G-J) Quantification of the percentage of cells with fragmented Golgi (G), TGN46 item number (H), area (I), and relative expression level (J) in F. (K) Immunoblots of Huh7-ACE2 cells incubated with or without SARS-CoV-2 (MOI = 2) for 24 h for indicated proteins. (L-S) Quantification of the relative expression of GRASP55 (L), GRASP65 (M), TGN46 (N), CatD (O), EEA1 (P), LC3-II/LC3-I ratio (Q), GPP130 (R) and ERGIC53 (S). Data are shown as mean ± SEM from at least three independent experiments. Statistical analyses are performed using two-tailed Student’s t-test. *, p < 0.05; **, p < 0.01; ***, p < 0.001; n.s., not significant.

To further validate the immunofluorescence results, we infected Huh7-ACE2 and Vero E6 cells with SARS-CoV-2 and blotted for key Golgi structural and functional proteins as well as a number of proteins in endosomes, lysosomes, and autophagosomes. Consistent with our immunofluorescence results, GRASP55 down-regulation and TGN46 up-regulation induced by SARS-CoV-2 infection were observed in both cell lines (Figure 4K, L, and N; Figure S5K, L, and N), while GRASP65 expression was not altered (Figure 4M; Figure S5M). SARS-CoV-2 infection also increased the precursor and total CatD level and elevated the LC3-II/LC3-I ratio while showing no impact on the expression of EEA1, GPP130, and ERGIC-53 (Figure 4O-S). Given that the infection rate was well below 100%, the actual changes were expected to be stronger than observed on the western blots. Taken together, GRASP55 was down-regulated and TGN46 was up-regulated by SARS-CoV-2 infection, suggesting that GRASP55 and TGN46 could serve as potential targets through which SARS-CoV-2 modulates the Golgi structure and function.

### GRASP55 expression significantly reduces SARS-CoV-2 infectivity and secretion

To test whether GRASP55 is a target of SARS-CoV-2 infection, we prepared stable Huh7-ACE2 cells expressing GFP or GFP-tagged wild-type (WT) GRASP55 by lentiviral transduction. Interestingly, cells that express GRASP55 displayed a significantly lower viral infection rate compared to cells that express GFP (Figure 5A-B). The spike pseudovirus entry assay showed that GRASP55 expression does not affect viral entry (Figure 5C and Figure S6A). To investigate the role of GRASP55 in SARS-CoV-2 trafficking, both cell lysates and culture media from infected GFP- or GRASP55-expressing cells were analyzed by western blotting. Both spike and N protein levels were decreased by about the same fraction in the lysate of GRASP55-expressing cells compared to that in GFP-expressing cells (Figure 5D, left panel, and Figure 5E-G), which is consistent with the decrease in viral infection demonstrated by IF (Figure 5A-B). While the spike and N protein levels were significantly reduced in the medium of GRASP55-expressing cells, the spike/N ratio was also significantly lower than that from GFP-expressing cells (Figure 5D, right panel, and Figure 5H-J), suggesting that GRASP55 expression may inhibit spike incorporation into virions. Further, both TCID50 and plaque assay show that viruses secreted from GRASP55-expressing cells possess significantly lower viral titer (Figure 5K-L).

**Figure 5.**
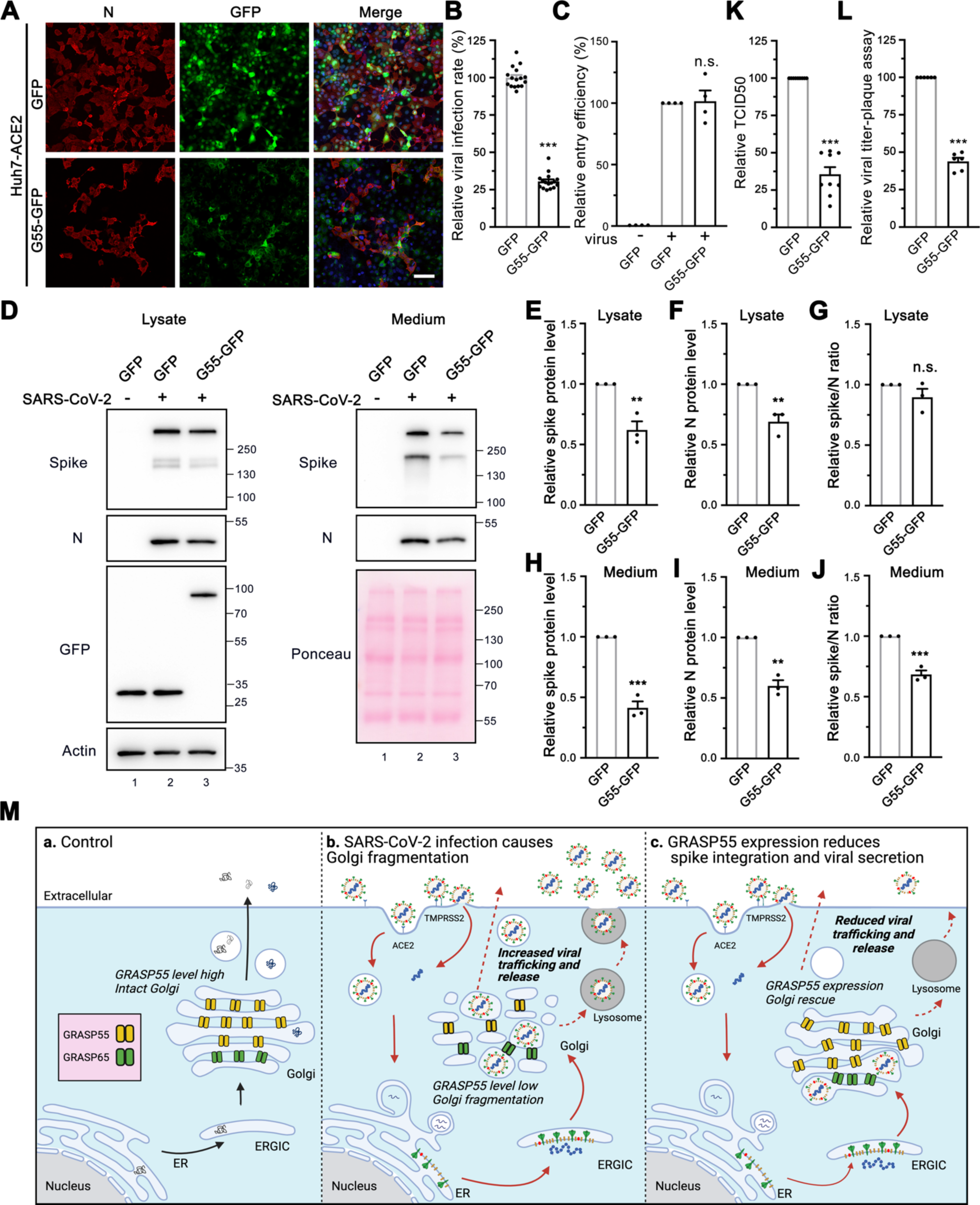
Expression of GRASP55 reduces SARS-CoV-2 assembly and secretion. (A) GRASP55 expression reduces SARS-CoV-2 infection. Stable Huh7-ACE2 cells expressing GFP or GRASP55-GFP were infected with SARS-CoV-2 (MOI = 3) for 24 h and stained for nucleocapsid. Shown are representative fluorescence images from 15 random images of one representative replicate from three independent experiments. Scale bars, 100 μm. (B) Quantification of the viral infection rate in A. (C) Cell entry assay of stable 293T-ACE2 cells expressing either GFP or GRASP55-GFP by SARS-CoV-2 Spike pseudotyped lentivirus. (D) Immunoblots of cell lysates and PEG-precipitated culture media of stable Huh7-ACE2 cells expressing GFP or GRASP55-GFP infected with SARS-CoV-2 (MOI = 3) for 24 h for indicated proteins. (E-J) Quantification of the relative expression of spike (E, H), N (F, I), and relative spike/N ratio (G, J) from cell lysates and culture media, respectively. (K-L) TCID50 assay (K) and plaque assay (L) of viruses collected from stable Huh7-ACE2 cells expressing GFP or GRASP55-GFP after SARS-CoV-2 infection (MOI = 3) for 24 h. (M) Proposed working model for a role of GRASP55 in SARS-CoV-2 infection. In brief, under normal conditions (a) GRASP55 is expressed at a high level and maintains the Golgi in an intact structure. After SARS-CoV-2 infection (b), GRASP55 level is reduced, resulting in Golgi fragmentation that may facilitate viral assembly and trafficking. Conversely, when GRASP55 is exogenously expressed (c), the Golgi structure is reinforced and the viral assembly and trafficking speed is limited, thus inhibiting spike incorporation and viral release. Data are shown as mean ± SEM. Statistical analyses are performed using two-tailed Student’s t-test. **, p < 0.01; ***, p < 0.001; n.s., not significant.

Previously we have shown that disruption of the stacked Golgi structure by depletion of GRASP55 and GRASP65 accelerates intra-Golgi trafficking possibly by increasing the membrane surface for vesicle formation (Xiang et al., 2013). This suggests a possibility that SARS-CoV-2 may utilize a similar mechanism to facilitate viral trafficking and release. To test whether GRASP55 expression affects spike protein trafficking, we co-expressed GRASP55 or GFP with spike and performed a cell surface biotinylation assay to compare the amount of spike protein at the plasma membrane. Much less spike was detected at the cell surface in GRASP55-expressing cells compared to GFP-expressing cells (Figure S6B, lane 8 vs. 7), although the total spike expression levels were comparable. This strongly suggests that GRASP55 expression hinders spike protein trafficking. Consistently, GRASP55 expression also reduced transferrin receptor (TfR) protein level at the cell surface, while this effect was enhanced by spike expression (Figure S6B). Further, we observed that Golgi fragmentation induced by SARS-CoV-2 infection was at least partially rescued by GRASP55 expression at all three infection time points (Figure S6C).

Based on these results, we propose a working model to explain the novel role of GRASP55 in SARS-CoV-2 trafficking (Figure 5M). Under normal conditions, GRASP55 is highly expressed, and the Golgi apparatus forms a stacked structure. SARS-CoV-2 infection decreases GRASP55 expression, resulting in Golgi fragmentation that may facilitate viral trafficking. When GRASP55 is exogenously expressed, the SARS-CoV-2 assembly and trafficking is reduced, thus limiting viral infectivity and secretion.

### GRASP55 depletion significantly enhances SARS-CoV-2 infectivity and secretion

To ensure that the inhibition of spike incorporation into virions and viral secretion caused by GRASP55 expression was not a result of global disruption of cell activities, we performed GRASP55 depletion assays. Both spike and N proteins were significantly increased, with a slight increase in the spike/N ratio in the lysate of GRASP55-depleted cells compared to that in siControl transfected cells (Figure 6A, left panel, and Figure 6B-D). Interestingly, the spike and N protein levels from the medium of GRASP55-depleted cells were increased by 2.4-fold and 1.6-fold, respectively, and the spike/N ratio was about 1.5-fold higher than that from siControl cells (Figure 6A, right panel, and Figure 6E-G), suggesting that GRASP55 depletion facilitates SARS-CoV-2 trafficking and spike incorporation. To investigate the impact of GRASP55 depletion on viral infection over time, we conducted a viral infection assay at three infection time points, ranging from 5 h to 24 h post-infection (about one to three viral replication cycles). There was no significant difference on the viral infection rate between control and GRASP55-depleted cells at 5 h and 8 h infection (Figure S7A-B), while at 24 h, the viral infection rate was about 1.5-fold higher in GRASP55-depleted cells compared to control cells (Figure S7A-B), which is consistent with the western blotting results (Figure 6A).

**Figure 6.**
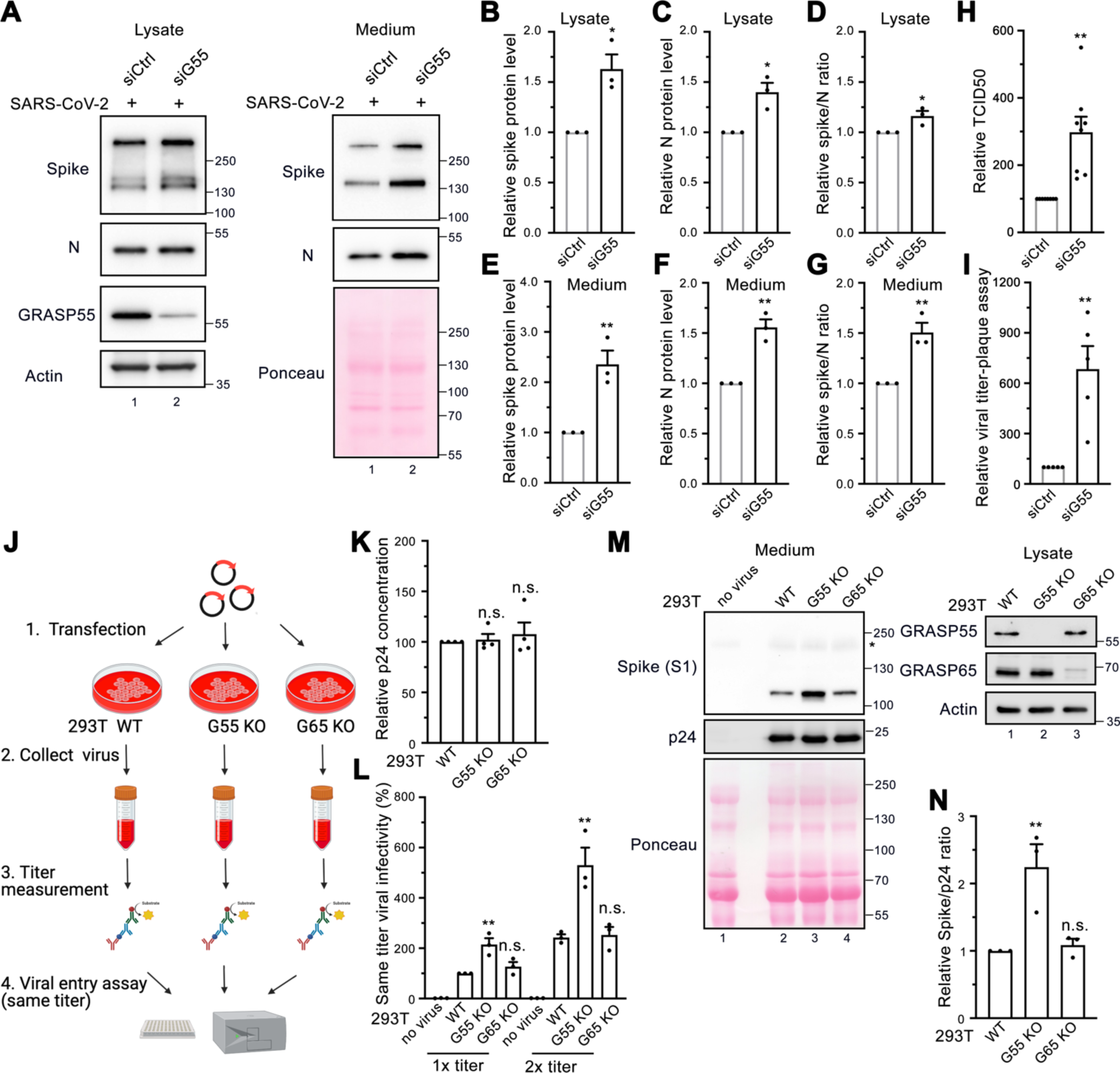
Depletion of GRASP55 accelerates SARS-CoV-2 assembly and secretion. (A) Immunoblots of cell lysates and PEG-precipitated culture media of Huh7-ACE2 transfected with siControl or siGRASP55 oligos for 48 h followed by SARS-CoV-2 infection (MOI = 3) for 24 h for indicated proteins. (B-G) Quantification of the relative expression of spike (B, E), N (C, F), and relative spike/N ratio (D, G) from cell lysates and culture media, respectively. (H-I) TCID50 assay (H) and plaque assay (I) of viruses collected from Huh7-ACE2 transfected with siControl or siGRASP55 oligos for 48 h followed by infection with SARS-CoV-2 (MOI = 3) for 24 h. (J-L) Diagram of a simplified protocol (J) of p24 titer measurement (K) and same-titer viral infectivity assay ( L) of two different titers from SARS-CoV-2 Spike pseudotyped lentiviruses collected from wild-type (WT), GRASP55 KO, and GRASP65 KO 293T cells. (M) Immunoblots of virus pellets prepared from wild-type, GRASP55 KO, and GRASP65 KO 293T cells for indicated proteins. GRASP55 and GRASP65 proteins are blotted from cell lysates on the right panel. (N) Quantification of the relative spike/p24 ratio of same-titer virus pellets prepared from wild-type, GRASP55 KO, and GRASP65 KO 293T cells. Data are shown as mean ± SEM. Statistical analyses are performed using two-tailed Student’s t-test (B-I) and one-way ANOVA, Tukey’s multiple comparison test (K-L, N), respectively. *, p < 0.05; **, p < 0.01; n.s., not significant.

Taking advantage of the readily accessible spike pseudotyped lentivirus in BSL2, we compared the secretion and infectivity of spike pseudotyped lentiviruses in WT and GRASP55 KO 293T cells (Figure 6J). Since our spike pseudotyped lentiviruses are HIV-based, the p24 level represents viral titer. Interestingly, spike pseudotyped lentiviruses prepared from GRASP55 KO cells at the same titer displayed a 2-fold higher viral infectivity compared to those from WT 293T cells. In contrast, GRASP65 KO had no effect on viral infectivity. To analyze the reasons for the difference in viral infectivity, ultracentrifugation was performed to concentrate the lentiviruses from three cell lines and viral pellets were analyzed by western blotting (Figure S7C). When the p24 level was used to normalize the virus amount in all three samples, the spike protein level in viruses produced by GRASP55 KO cells was dramatically higher than that by WT cells and GRASP65 KO cells (Figure 6M). The spike/p24 ratio was more than 2-fold higher for GRASP55 KO cells produced viruses compared to WT cells (Figure 6N), suggesting that GRASP55 KO accelerates spike trafficking to the cell surface where lentiviruses are assembled, thus leading to a higher number of spikes on the viral particles. Altogether, GRASP55 depletion significantly promotes spike trafficking through the secretory pathway to deliver more spike protein to sites of SARS-CoV-2 assembly.

### TGN46 is required for viral trafficking of both WA-1 and Delta variants of SARS-CoV-2

The finding that TGN46 protein level is significantly increased in SARS-CoV-2 infected cells indicates a need for TGN46 in viral infection. Therefore, we speculated that depletion of TGN46 might most likely inhibit SARS-CoV-2 infection. Indeed, TGN46 depletion significantly inhibited infection by the SARS-CoV-2 USA-WA1 strain (Figure 7A-B). To determine if this effect is specific for the WA1 strain, we also tested the Delta variant. Similar to the WA1 strain, TGN46 depletion also reduced infection by the Delta variant (Figure 7C-D). These results demonstrate a role for TGN46 in SARS-CoV-2 infection independent of the viral strain.

**Figure 7.**
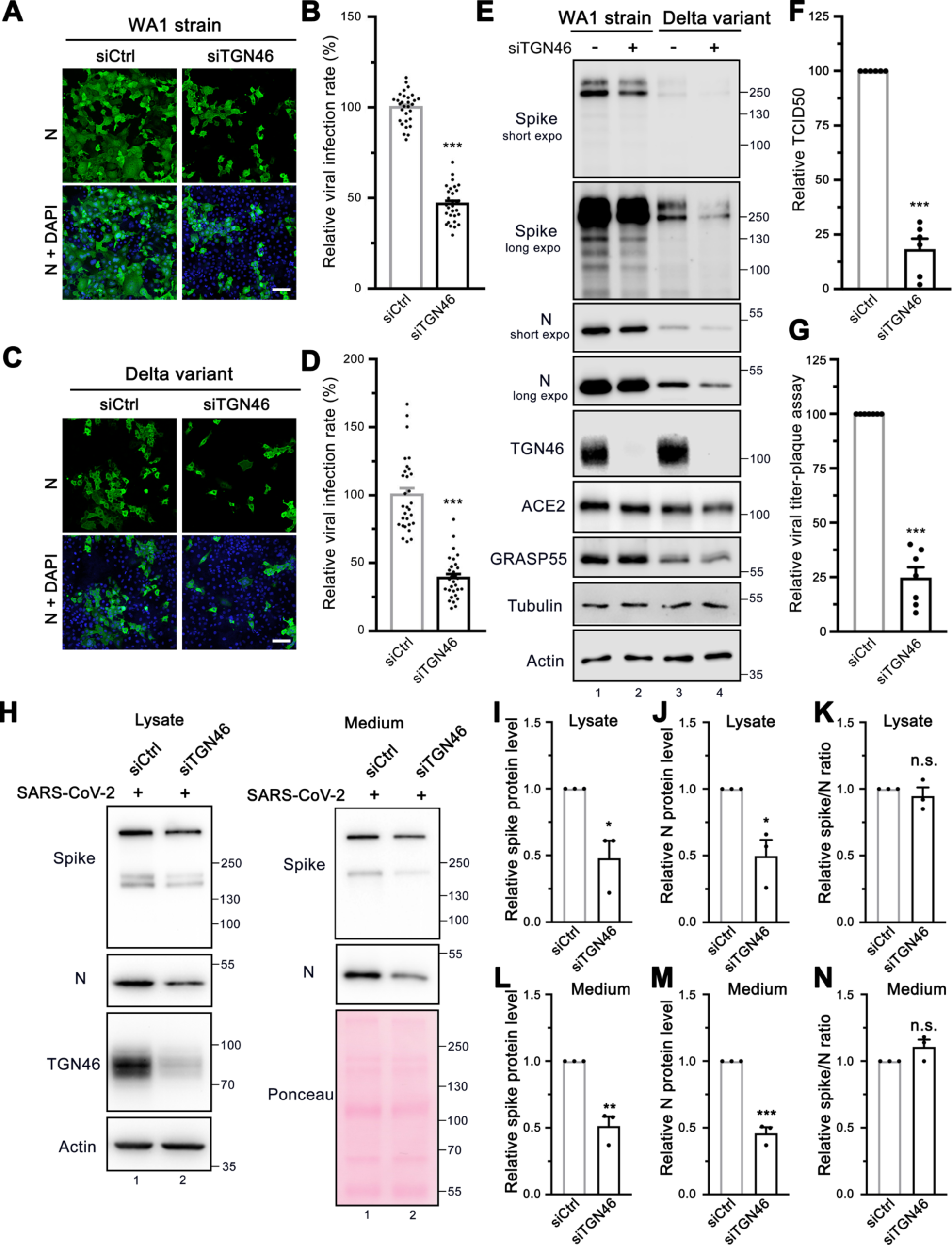
TGN46 is required for SARS-CoV-2 trafficking. (A, C) TGN46 depletion reduces SARS-CoV-2 infection. Huh7-ACE2 cells were transfected with siControl or siTGN46 oligos for 48 h followed by infection (MOI = 1) with the WA1 strain (A) or Delta variant (C) of SARS-CoV-2 for 24 h and stained for nucleocapsid. Shown are representative fluorescence images from 30 random images from two independent experiments. Scale bars, 100 μm. (B, D) Quantification of the viral infection rate in A and C, respectively. (E) TGN46 depletion reduces viral protein expression in host cells. Huh7-ACE2 cells were transfected with siControl or siTGN46 oligos for 48 h followed by infection with SARS-CoV-2 WA1 strain and Delta variant (MOI = 3) for 24 h. Cell lysates were blotted for indicated proteins. Long and short exposures are shown for spike and N proteins. (F-G) TCID50 assay (F) and plaque assay (G) of viruses collected from Huh7-ACE2 transfected with siControl or siTGN46 oligos for 48 h followed by infection with SARS-CoV-2 for 24 h. (H) Immunoblots of cell lysates and PEG-precipitated culture media of Huh7-ACE2 transfected with siControl or siTGN46 oligos for 48 h followed by SARS-CoV-2 infection (MOI = 3) for 24 h for indicated proteins. (I-N) Quantification of the relative expression of spike (I, L), N (J, M), and relative spike/N ratio (K, N) from cell lysates and culture media, respectively. Data are shown as mean ± SEM. Statistical analyses were performed using two-tailed Student’s t-test. *, p < 0.05; **, p < 0.01; ***, p < 0.001; n.s., not significant.

Consistent with the viral infection assays, depletion of TGN46 reduced the expression of both spike and nucleocapsid viral proteins when cells were infected with either the USA-WA1 strain or the Delta variant, confirming a critical role for TGN46 in SARS-CoV-2 infection with both strains (Figure 7E). TGN46 depletion has no effect on viral entry (Figure S8A-B) but reduced viral production, as shown by both TCID50 and plaque assays (Figure 7F-G), suggesting that TGN46 depletion inhibits SARS-CoV-2 secretion.

Given that TGN46 is recycling between the Golgi and plasma membrane, we speculated that it may facilitate viral trafficking from the Golgi to the plasma membrane. To test the role of TGN46 in spike trafficking, we transfected control and TGN46-depleted Huh7-ACE2 cells with spike and performed cell surface biotinylation and streptavidin pulldown. The amount of spike protein at the cell surface was dramatically decreased after TGN46 depletion, which was revealed by both strep and spike antibodies (Figure S8C, lane 8 vs. 7). In spike-expressing cells, not only was spike affected by TGN46 knockdown, but several other cell surface proteins, including ACE2, insulin-like growth factor 2 receptor (IGF2R), TfR, and E-cadherin, also displayed a reduction at the plasma membrane in TGN46 knockdown cells (Figure S8C, lane 8 vs. 7). The effect of TGN46 knockdown on ACE2, IGF2R and TfR cell surface localization was not detected in non-spike expressing cells (Figure S8C, lane 6 vs. 5), implying that spike expression or viral infection may hijack the majority of TGN46 for viral trafficking and so post-Golgi trafficking of many cell receptors to the plasma membrane is impeded. Notably, the level of TGN46 at the cell surface was increased after spike expression (Figure S8C, lane 7 vs. 5), indicating that TGN46 may recycle more frequently between the TGN and the plasma membrane when spike is expressed.

To determine if TGN46 functions in a similar manner to GRASP55 in the regulation of both viral assembly and trafficking, both cell lysate and medium were analyzed by western blotting. Both spike and N were decreased in cell lysate and medium of TGN46-depleted cells (Figure 7H-J and L-M), similar to GRASP55 expression. However, the spike/N ratio remained unchanged in both lysate and medium after TGN46 depletion (Figure 7K and 7N), suggesting that TGN46 modulates SARS-CoV-2 trafficking but not assembly and that TGN46 and GRASP55 function differently. Taken together, we propose a working model of TGN46 in SARS-CoV-2 trafficking (Figure S8D). In brief, SARS-CoV-2 infection significantly increases TGN46 protein level to accelerate viral trafficking without affecting viral assembly and entry, while TGN46 depletion inhibits the trafficking and release of all variants of SARS-CoV-2. Our study revealed TGN46 as a promising target for COVID-19 treatment.

### GRASP55 and TGN46 depletion additively inhibits viral trafficking

To determine whether GRASP55 expression and TGN46 depletion display an additive effect on inhibiting SARS-CoV-2 trafficking, we depleted TGN46 in stable GFP- or GRASP55-GFP expressing cells. Consistent with previous results, expression of GRASP55 or depletion of TGN46 separately decreased spike and N protein levels in both cell lysate and medium (Figure 8A, lane 1 vs lane2, lane 1 vs lane 3, Figure 8B-C and 8E-F). We observed a dramatic reduction of both spike and N in cell lysate and medium from GRASP55-expressing and TGN46-depleted cells, which was greater in medium than that in lysate (Figure 8A-C and 8E-F). The spike/N ratio was not altered in the lysate of the four cell lines (Figure 8D), however, GRASP55 expression in either siControl transfected cells or siTGN46-transfected cells significantly decreased the spike/N ratio in the medium (Figure 8G). A combination of GRASP55 expression and TGN46 depletion also displayed greater inhibition in viral secretion than either alone (Figure 8H-I). Our results demonstrate that GRASP55 regulates both viral assembly and trafficking, while TGN46 only modulates viral trafficking (Figure 8J).

**Figure 8.**
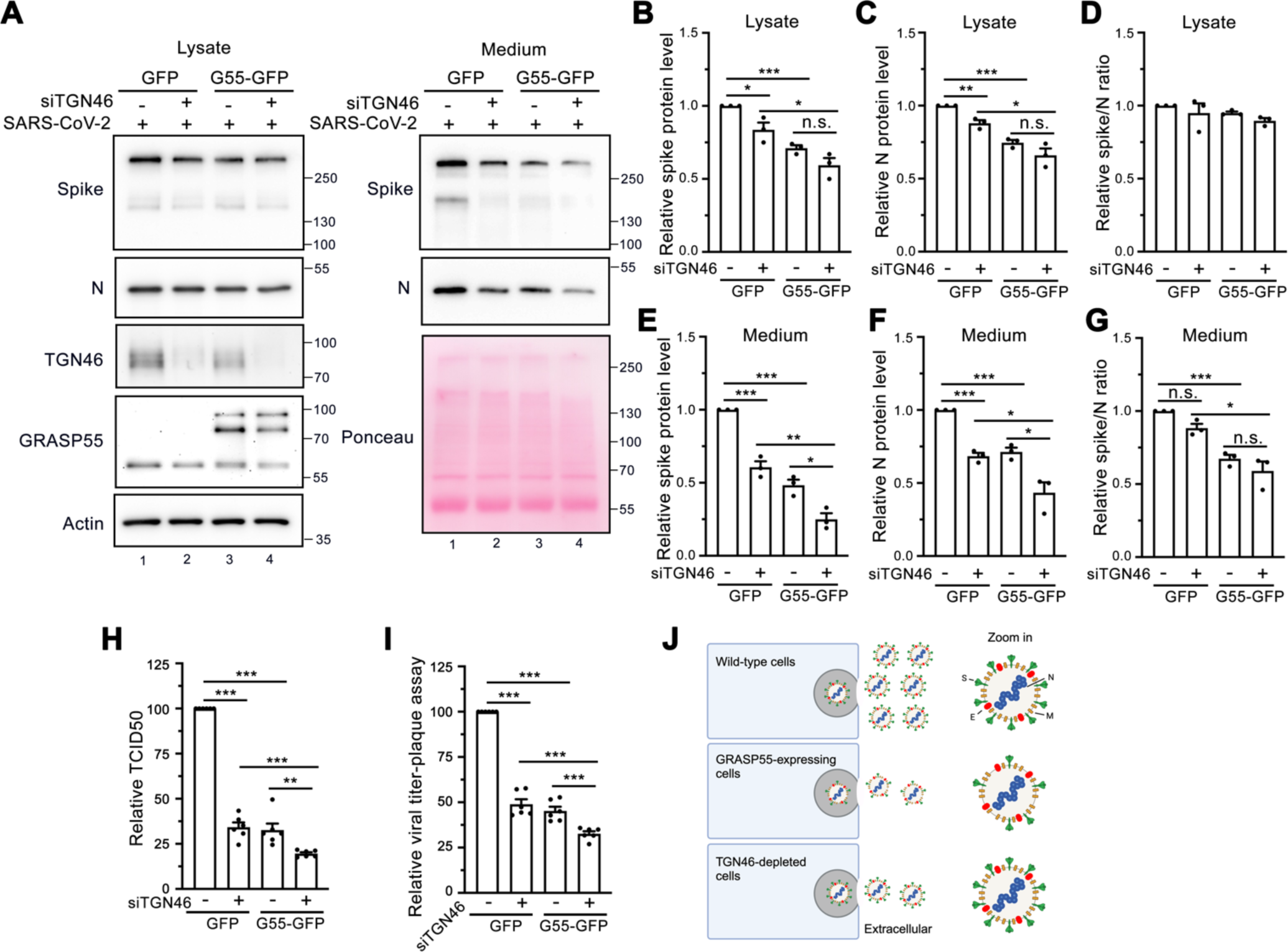
Expression of GRASP55 and depletion of TGN46 additively inhibit viral trafficking. (A) Immunoblots of cell lysates and PEG-precipitated culture media of Huh7-ACE2 cells stably expressing GFP or GRASP55-GFP and transfected with either siControl or siTGN46 oligos for 48 h followed by SARS-CoV-2 infection (MOI = 3) for 24 h for indicated proteins. (B-G) Quantification of the relative expression of spike (B, E), N (C, F), and the relative spike/N ratio (D, G) from cell lysates and culture media, respectively. (H-I) TCID50 assay (H) and plaque assay (I) of viruses collected from Huh7-ACE2 cells stably expressing GFP or GRASP55-GFP and transfected with either siControl or siTGN46 oligos with SARS-CoV-2 for 24 h. (J) A simplified working model illustrating the effects of GRASP55 and TGN46 expression levels on SARS-CoV-2 assembly and secretion. In brief, GRASP55 expression inhibits both SARS-CoV-2 spike incorporation into virions and viral trafficking, while TGN46 depletion only affects viral trafficking, with no impact on viral assembly. Data are shown as mean ± SEM. Statistical analyses were performed using two-tailed Student’s t-test. *, p < 0.05; **, p < 0.01; ***, p < 0.001; n.s., not significant.

## Discussion

In this study, we revealed that SARS-CoV-2 infection triggers a global change of the endomembrane system of host cells, particularly affecting the Golgi structure. Conversely, Golgi disruption by BFA strongly inhibits viral infection but not viral entry. To determine the mechanism of Golgi fragmentation induced by SARS-CoV-2 infection, we surveyed a large number of Golgi proteins in SARS-CoV-2 infected cells. While several Golgi proteins are impacted by viral infection, GRASP55 and TGN46 are the top two proteins whose levels change most dramatically in opposite trends. Surprisingly, overexpression of GRASP55 not only decelerates viral trafficking but also inhibits viral assembly, while GRASP55 depletion reverses the effects. Distinct from GRASP55, TGN46 plays a role in SARS-CoV-2 trafficking but not its assembly. Manipulation of either GRASP55 or TGN46 does not affect viral entry. Finally, we show that GRASP55 and TGN46 modulate SARS-CoV-2 trafficking at different stages and display an additive effect. Thus, our study uncovers a novel mechanism by which SARS-CoV-2 modulates the Golgi structure via regulating GRASP55 and TGN46 expression to facilitate viral assembly and trafficking. Our results indicate that the Golgi apparatus may serve as a novel therapeutic target for the treatment of COVID-19 and other diseases caused by viruses that utilize a similar trafficking pathway.

The Golgi stacking proteins, GRASP55 and GRASP65, play essential roles in Golgi structure formation (Tang and Wang, 2013; Xiang and Wang, 2010) by forming trans-oligomers that “glue” adjacent Golgi cisternae together into stacks and ribbon (Tang et al., 2012; Tang et al., 2010b). Expression of phospho-deficient mutants of GRASPs (e.g., the GRASP domain of GRASP55 or GRASP55) at least partially inhibits mitotic Golgi disassembly (Wang et al., 2005; Xiang and Wang, 2010). Inhibition of Golgi stacking in cells by microinjecting GRASP antibodies or by GRASP55/65 knockdown/knockout (KD/KO) (Tang et al., 2010b; Wang et al., 2003) accelerates conventional protein trafficking but impairs accurate glycosylation and sorting (Bekier et al., 2017; Wang et al., 2008; Xiang et al., 2013), increases heparan sulfate but decreases chondroitin sulfate synthesis (Ahat et al., 2022b), and reduces unconventional protein secretion of mutant huntingtin (Ahat et al., 2022a). A plausible explanation is that stacking may reduce the accessibility of coat proteins to Golgi membranes, which decreases the rate of vesicle budding and fusion (Huang and Wang, 2017; Zhang and Wang, 2015, 2016). Therefore, it is reasonable to speculate that Golgi fragmentation observed in this study may facilitate SARS-CoV-2 trafficking and release.

SARS-CoV-2 infection down-regulates GRASP55 but not its homolog GRASP65. GRASP55 has been shown to play a crucial role in various stress responses (Ireland et al., 2020; Nuchel et al., 2021; Zhang and Wang, 2018, 2020), while GRASP65 functions more in cell migration and apoptosis (Ahat et al., 2019; Bisel et al., 2008; Lane et al., 2002). In this study, we found that GRASP55 plays a vital role in viral assembly and secretion, likely due to its role in Golgi structure formation. Indeed, GRASP55 expression significantly inhibited spike protein trafficking to the cell surface, supporting our hypothesis that SARS-CoV-2 induced GRASP55 down-regulation facilitates viral trafficking and spike incorporation into virions, consistent with the previous finding that GRASP55-depletion induced Golgi structural defect enhances conventional protein trafficking (Xiang et al., 2013). It is believed that SARS-CoV-2 virions assemble within the ERGIC (V’Kovski et al., 2021) and Golgi (Cortese et al., 2020; Scherer et al., 2022). Therefore, it is reasonable that GRASP55 regulates SARS-CoV-2 spike incorporation into virions. Given the pivotal role of the spike protein in viral infection, the finding that GRASP55, a Golgi structural protein, regulates both viral production and infectivity is intriguing. It would be compelling to determine whether M or E protein is also increased on each virion by GRASP55 depletion.

Many enveloped and non-enveloped viruses display trimeric receptor-binding proteins at the viral surface (e.g. SARS-CoV-2 spike, influenza HA) that are usually about 10 nm or longer to play a crucial role in mediating cell entry (Dimitrov, 2004). The number of these receptor-binding proteins per virion is a determinant of viral infectivity. Among enveloped RNA viruses, this number varies from 7-14 (HIV), 60 (Zika virus), 400-500 (influenza virus), to 1200 (vesicular stomatitis virus). Three independent teams reported that there were on average 24-40 spikes on each SARS-CoV-2 virion (Ke et al., 2020; Turonova et al., 2020; Yao et al., 2020). In our study, GRASP55 depletion increased the spike number on secreted SARS-CoV-2 virions about 1.5-fold. This increase in the spike number significantly enhanced viral infectivity (Figure 6H-I), and it is likely to be tolerated by virions, given that influenza virus has 400-500 HA trimers per virion and a diameter (about 100 nm in the spherical form) similar to SARS-CoV-2 (100-120 nm). Whether GRASP55 depletion affects SARS-CoV-2 viral particle size remains an open and interesting question.

TGN46 is a glycoprotein that recycles between the TGN and cell surface (Ponnambalam et al., 1994; Reaves et al., 1993). TGN46 was previously reported to directly interact with integrin β1 to modulate its trafficking (Wang and Howell, 2000), while recently TGN46 was reported to serve as a sorting receptor at the TGN for CARTS, a class of protein kinase D-dependent TGN-to-plasma membrane carriers (Lujan et al., 2022). Therefore, we postulate that the increased TGN46 level indicates a need for TGN46 to accommodate the high flux of virion trafficking through the late secretory pathway, and our results support this hypothesis. Notably, depletion of TGN46 inhibited spike protein trafficking, but not the amount of ACE2 at the cell surface, while spike expression increased the TGN46 level at the cell surface. These observations indicate that TGN46 functions in the post-Golgi trafficking of the SARS-CoV-2 virus. Our discoveries that SARS-CoV-2 reduces GRASP55 and elevates TGN46 expression are also supported by a number of RNA-seq and proteomic studies of SARS-CoV-2 infected cells and human tissues (Blanco-Melo et al., 2020; Bojkova et al., 2020; Sun et al., 2020). Significantly, our results demonstrate that either overexpression of GRASP55 or depletion of TGN46 inhibits SARS-CoV-2 secretion.

It is noteworthy that both Tg and BFA inhibit viral infection rate but promote viral entry. It was reported that BFA treatment for a short period (4-6 h) did not affect the mouse hepatitis virus (MHV) egress (Ghosh et al., 2020), while our BFA treatment for 24 h dramatically inhibited SARS-CoV-2 infection, suggesting that integrity of the conventional trafficking pathway is required for SARS-CoV-2 replication. Although it has been shown that CQ does not block SARS-CoV-2 infection in the human lung cell line Calu-3 (Hoffmann et al., 2020b), it greatly reduces both viral infection rate and viral entry efficiency in Huh7-ACE2 cells in our study. This discrepancy may be explained by the fact that SARS-CoV-2 entry into Calu-3 cells is independent of endosomal acidification due to its high expression level of TMPRSS2, which activates spike at the plasma membrane; while Huh7.5 (a derivative cell line of Huh7) cells heavily rely on the endosomal acidity for SARS-CoV-2 entry (Dittmar et al., 2021), leading to a high dependence on lysosomes whose function can be inhibited by CQ. Vacuolin-1 is sometimes regarded as a lysosomal exocytosis inhibitor, but it has also been reported to alter lysosome morphology without inhibiting Ca^2+^-regulated lysosomal exocytosis (Huynh and Andrews, 2005). Although we showed that vacuolin-1 inhibits viral entry, it is unclear whether the inhibition of SARS-CoV-2 infection by vacuolin-1 observed in our study is also contributed by the inhibition of SARS-CoV-2 release via lysosomal exocytosis.

Currently, it is still debated about whether SARS-CoV-2 virus transits the Golgi during secretion (Sergio et al., 2024). However, our study strongly supports that Golgi is required for SARS-CoV-2 secretion for the following reasons. Firstly, we observed that many mature SARS-CoV-2 virions reside in both rims of a stacked Golgi and fragmented Golgi remnants by electron microscopy. Secondly, two Golgi proteins, GRASP55 whose level is closely related to the Golgi structure, and TGN46 which constitutively recycles from TGN and plasma membrane, play important roles in SARS-CoV-2 trafficking, suggesting that the Golgi is a key target utilized by SARS-CoV-2. Lastly, our findings that GRASP55 protein regulates SARS-CoV-2 spike number on each virion indicate that the Golgi affects spike incorporation into virions and thus viral infectivity. In summary, our study reveals that the Golgi plays a key role in SARS-CoV-2 assembly and secretion, and it could be a potential target for blocking the spread of different SARS-CoV-2 variants.

## Methods and Materials

### Generation of cDNA constructs

All pCAG plasmids encoding Strep-tagged SARS-CoV-2 proteins were kind gifts from Dr. Nevan Krogan (Gordon et al., 2020). pHAGE2-pEF1α-GRASP55-GFP was constructed by cloning GRASP55-GFP from pEGFP-N1-GRASP55 into the pHAGE2-pEF1α vector. psPAX2 and pMD2G were obtained from Addgene. A C-terminal 19 aa truncated SARS-CoV-2 spike expression vector and CMV-eGF1 were kind gifts from Dr. Marilia Cascalho.

### Cell culture and transfection

ACE2-expressing Huh7 cells were sorted, enriched, and selected by flow cytometry (FACS) to ensure a high infection rate of SARS-CoV-2 (Sherman et al., 2021). Huh7-ACE2 and Vero E6 (ATCC CRL 1586) cells were cultured in Dulbecco’s Modified Eagle’s Medium (Gibco) supplemented with 10% fetal bovine serum (FBS, Hyclone), 100 units/ml penicillin and 100 μg/ml streptomycin (Invitrogen) at 37°C with 5% CO2. Plasmid transfection was performed using Lipofectamine 2000 or Lipofectamine 3000 (Invitrogen) following the manufacturer’s instructions. After 24-48 h, transfected cells were fixed by 4% paraformaldehyde (PFA) for immunofluorescence analysis or were lysed in IGEPAL-C360 lysis buffer (20 mM Tris HCl pH 8.1, 37 mM NaCl, 1% IGEPAL-C360, 10% glycerol, 2 mM EDTA supplemented with protease inhibitor cocktail from ThermoFisher) for immunoblotting.

### SARS-CoV-2 viral strains and amplification

Most experiments were conducted using SARS-CoV-2, Isolate USA-WA1/2020 (BEI NR-52281), unless specified. The Delta variant of SARS-CoV-2 used in the study is the Isolate hCoV-19/USA/PHC658/2021 (Lineage B.1.617.2; BEI NR-55611). SARS-CoV-2 working stocks were amplified by infecting Vero E6 cells in 2% FBS DMEM and 182 cm^2^ flasks with 70-80% confluency. Flasks were incubated until cytopathic effect (CPE) became distinctly visible, generally at 2-4 days post-infection (dpi). The cell debris was pelleted and removed by centrifugation, and the supernatant was harvested, filtered through 0.45 µm SCFA syringe filters and pipetted into 800 µl aliquots, with tubes of viral stock then being stored at −80°C. One vial of each propagated stock was thawed after 24 h and titer was measured by TCID50, with positive wells determined by the presence of CPE 4 dpi observed by 4x objective phase-contrast microscopy. Viral titer was calculated by the modified Reed and Muench method. All virus inactivation protocols were validated and approved by the Institutional Biosafety Committee at University of Michigan.

### RNA interference

siRNA Universal Negative Control #1 (siControl), siGRASP55 with 5’-UGAUAACUGUCGAGAAGUGAUUAUU-3’ sequence, siGRASP65 with 5’-CCUGAAGGCACUACUGAAAGCCAAU-3’ sequence, and siTGN46 with 5’-CCACCGAAAGCGUCAAGCAAGAAGA-3’ sequence were obtained from Sigma-Aldrich. Huh7-ACE2 cells were transfected with siRNA oligos (final concentration, 50 nM) by RNAi-MAX (Invitrogen) on day 1 and infected with SARS-CoV-2 on day 3. Cells were fixed by 4% paraformaldehyde (PFA) for 30 min on day 4 for immunofluorescence analysis.

### Immunoblotting

Huh7-ACE2 or Vero E6 cells were seeded onto 6-well plates on day 1 and incubated with or without SARS-CoV-2 (MOI = 2) on day 2 for 24 h. Cells were lysed in IGEPAL-C360 lysis buffer. For regular immunoblotting of uninfected cells, cells were lysed in lysis buffer (20 mM Tris-HCl pH 8.0, 150 mM NaCl, 1% Triton X-100 supplemented with protease inhibitor cocktail (Thermo Fisher). The homogenate was centrifuged at 13000 × *g* for 15 min to remove cell debris, then denatured at 95°C for 10 min in 2x Laemmli buffer supplemented with 5% 2-mercaptoethanol. Protein quantification was performed using a Bradford Kit (Bio-Rad). Protein samples were analyzed by SDS-PAGE and then transferred to nitrocellulose membranes using a semi-dry or wet transfer machine. The membranes were blocked in 5% milk in PBST (0.1% Tween 20 in phosphate buffered saline) and incubated with proper antibodies and visualized by a FluorChem M chemi-luminescent imager (ProteinSimple, San Jose, CA) with enhanced chemiluminescence (Thermo Fisher). The antibodies used in this study are shown in Supplementary Table 1.

### Immunofluorescence

Huh7-ACE2 or Vero E6 cells were seeded on poly-lysine-coated coverslips. SARS-CoV-2 infected cells were fixed with 4% PFA for 30 min for complete virus inactivation. Cells were quenched with 50 mM NH4Cl in PBS for 10 min with gentle rocking and permeabilized with 0.2% Triton X-100 for 10 min. Cells were blocked with 0.2% gelatin blocking buffer in PBS (PGA) for 30 min at room temperature, incubated with primary antibodies overnight and secondary antibodies for 1 h. The antibodies used for immunofluorescence were shown on Supplementary Table 1. Hoechst 33258 (Sigma-Aldrich) was used to stain the nuclear DNA. About 15 to 20 random images were taken for each independent experiment with a 60x oil objective on a Nikon ECLIPSE Ti2 Confocal microscope and processed with maximum intensity projection. Quantifications were performed to calculate item number, area, and sum intensity of selected ROIs using the Nikon NIS-Elements AR analysis software. Control and virus-infected samples were processed in parallel following the same procedure. All images for the same marker were captured and processed with the same setting. For 3D video reconstruction, samples were prepared as above. Z-stacks of 40 images were taken in 0.15 µm increments. Maximum intensity projection was performed, and videos were made with the Nikon NIS-Elements AR analysis software.

### Viral infection rate assay

Huh7-ACE2 cells were seeded onto poly-lysine-coated coverslips on day 1. On day 2, cells were treated with or without indicated chemicals and immediately infected with SARS-CoV-2 for 24 h. Cells were fixed with 4% PFA for 30 min and processed for immunofluorescence analysis. Images were taken with a 20x air objective on a Nikon ECLIPSE Ti2 Confocal microscope and processed with maximum projection. Cell numbers for uninfected or infected cells were counted by the counting tool in Adobe Photoshop using nucleocapsid as a virus marker and the viral infection rate [infected cells/(uninfected + infected cells)] for DMSO treatment was normalized to 100%.

### TCID50 assay

Vero E6 cells were seeded into a 96-well plate at the seeding density of 1 x 10^4^ cells/well in 100 µl full culture medium. When cells were close to confluency, 125 µl pure DMEM medium without FBS was added to each well. 25 µl viral stock was added to the first column (8 wells) and mixed well. 25 µl mixture from the first column was taken and added into the second column. This step was repeated to obtain a serial dilution with 8 replicates for each sample and 7 dilutions. After incubation for 4-5 days, the cells were observed under microscope for each well and TCID50 was calculated.

### Plaque assay

Vero E6 cells were seeded into a 6-well plate at the seeding density of 3 x 10^5^ cells/well. When cells were close to confluency, culture medium was discarded and 1 ml DMEM medium with 2% FBS was added. 100 µl virus with a total of 5 serial dilution was added. Cells were incubated with viruses for 1 h and mixed by moving plates back and forth every 15 min. Infection medium was aspirated and 2 ml pre-warmed carboxymethylcellulose (CMC) overlay medium (DMEM medium with 2.5% FBS and 1% CMC) was added. Cells were incubated in the incubator for 4-5 days. CMC overlay medium was aspirated, and cells were incubated with 0.5 ml 4% PFA in PBS for 20 min at room temperature. PFA was removed before adding 0.1% crystal violet in 20% methanol for 20 min at room temperature. Cells were gently rinsed with water. Plaques are counted after drying and virus titer was calculated.

### PEG virus precipitation assay

PEG virus precipitation assay was performed to concentrate viruses from the culture medium of Huh7-ACE2 cells, by following the product instructions (Cat #MAK343, Sigma). In brief, after 24 h infection, cells were lysed using RIPA buffer, and the 10 ml media were centrifuged at 3,200 g for 15 min at 4°C to remove cell debris. Supernatants were transferred into new tubes and mixed with 2.5 ml PEG solution (5x) at 4°C overnight. After centrifugation at 3,200 g for 30 min, supernatant was carefully removed, and the white pellet was resuspended by 100 µl virus resuspension solution and added with RIPA buffer, then denatured at 95°C for 10 min in 2x Laemmli buffer supplemented with 5% 2-mercaptoethanol for WB analysis.

### Stable cell lines preparation by lentivirus

HEK293T cells were seeded into 10-cm dishes at a seeding density of 300,000 cells/ml on day 1. On day 2, cells were co-transfected with three plasmids, pHAGE2-pEF1α-GFP or pHAGE2-pEF1α-GRASP55-GFP, psPAX2, and pMD2.G, by Lipofectamine 3000. Culture medium was changed 6 h post transfection. Lentiviruses were collected after 24 h and 48 h transfection, filtered using PAS filter with a 0.45 µm pore size, and stored at −80°C. Huh7-ACE2 or 293T-ACE2 cells were seeded into 6-cm dishes. Next day, cells were infected by lentiviruses expressing GFP or GRASP55-GFP with 10 µg/ml polybrene. Puromycin at a final concentration of 2 µg/ml was added into cells 48 h post infection. Transfection efficiency was validated by immunofluorescence after continuous puromycin treatment for 3-4 days.

### Spike pseudotyped lentivirus preparation and titer measurement

Spike pseudotyped lentiviruses were prepared using 293T cells co-transfected with pLenti-pseudo spike (a SARS-CoV-2 spike expression vector), psPAX2, and CMV-eFP1 by Lipofectamine 3000. All other steps are the same with GRASP55-GFP expressing lentivirus preparation as described above. Virus titer was measured by using Lenti-X p24 Rapid Titer Kit (Cat #022261, Takara).

### Spike pseudotyped lentivirus ultracentrifugation

Lentiviruses were collected after 24 h and 48 h transfection, filtered using PAS filter with a 0.45 µm pore size, and titered by Lenti-X p24 Rapid Titer Kit. Same titered viruses were prepared by adding fresh culture medium into the same volume. Ultracentrifugation was performed using a SW-41 Ti rotor, at 30,000 rpm (110,682 g) for 2 hours at 4°C. The supernatant was discarded and virus pellet was resuspended in the same volume of RIPA buffer (Cat #89901, ThermoFisher, 20 mM Tris-HCl, pH 7.6, 150 mM NaCl, 1% NP-40, 0.1% SDS, 1% sodium deoxycholate, and 1× protease inhibitor cocktail) and sample buffer for further western blotting analysis.

### Virus cell entry assay

293T-ACE2 cells were seeded into 96-well plates (Corning #3903) at the seeding density of 300,000 cells/ml on day 1. On day 2, cells were infected by Spike pseudotyped lentiviruses with 8 µg/ml polybrene and were co-treated with small molecules or transfected with siControl or siTGN46 48 h prior to infection. On day 3, cells were lysed with the Bright Glo Kit reagent for 5 min followed by immediate reading using Tecan SparkCyto with the luminescence mode. All experiments were done in four replicates and repeated at least three times.

### Cell surface biotinylation assay

Cell surface biotinylation assay was performed as previously described (Ahat et al., 2019). All procedures were performed on ice or at 4°C. In brief, Huh7-ACE2 cells were seeded on 15-cm dishes. After knocking down TGN46 or overexpression of GRASP55, cells were washed twice with ice-cold PBS, treated with 10 ml of 0.5 mg/ml NHS-SS-biotin (Thermo Fisher) in PBS for 20 min in the cold room, and quenched by 100 mM glycine in PBS for 10 min. After three washes with ice-cold PBS, cells were lysed in lysis buffer (20 mM Tris-HCl, pH 8.0, 150 mM NaCl, 1% Triton X-100, 0.1% SDS, 0.1% sodium deoxycholate, 1 mM EDTA, 50 mM sodium fluoride, 20 mM sodium orthovanadate, and 1× protease inhibitor cocktail). After centrifugation, the supernatants were adjusted to the same concentration and incubated with streptavidin-agarose beads overnight. After extensive wash, beads were boiled in SDS loading buffer with 40 mM DTT. Proteins were separated by SDS-PAGE and analyzed by immunoblotting.

### Electron microscopy (EM)

All EM-related reagents were purchased from Electron Microscopy Sciences (EMS; Hatfield, PA). Huh7-ACE2 cells were infected with or without SARS-CoV-2 (MOI = 1) for 24 h and fixed in 2% glutaraldehyde at room temperature for 30 min. Cells were washed three times with 0.1 M sodium cacodylate, and post-fixed on ice in 1% (wt/vol) reduced osmium tetroxide, 0.1 M sodium cacodylate and 1.5% (wt/vol) cyanoferrate. Cells were rinsed 3 times with 0.1 M sodium cacodylate and 3 times with water, and processed for successive dehydration and embedding as previously described (Tang et al., 2010a). Resin blocks were cut to ultrathin (50-70 nm) sections with a diamond knife and mounted on Formvar-coated copper grids. Grids were double contrasted with 2% uranyl acetate for 5 min and then with lead citrate for 2 min and washed with excess water. Images were captured at 1,500x, 3,000x, 8,000x, and 12,000x magnifications by a JEOL JEM-1400 transmission electron microscope.

### Quantitation and statistics

All data represent the mean ± SEM of at least three independent experiments unless noted. Statistical analyses were performed by two-tailed Student’s t-test, or one-way ANOVA with Tukey’s multiple comparison test using GraphPad Prism9. Differences in means were considered as no statistical significance (n.s.) if p ≥ 0.05. Significance levels are as follows: *, p < 0.05; **, p < 0.01; ***, p < 0.001. Figures are assembled with Photoshop CS6 Extended (Adobe, San Jose, CA). Figures 5M, 8J and S7D are created with BioRender.

## ACKNOWLEDGMENTS

We thank Dr. Nevan Krogan for kind gifts of plasmids encoding Strep-tagged SARS-CoV-2 proteins; Drs. Zhe Han, Jin-Gu Lee, Weichao Zhang, Ming Li, Meiqin Hu, and Haoxing Xu for reagents; and Dr. Gregg Sobocinski for technical assistance on electron microscopy. We thank Drs. Christiane Wobus, Akira Ono, Lois Weisman, Kenneth Cadigan, Mohammed Akaaboune, Ming Li, and current and past members of the Wang lab, especially Erpan Ahat and Jie Li, for stimulating discussions and suggestions. A.W.T. was supported by the NIH (Grants R21AI152865 and R01GM139823), Y. W. was supported by the NIH (Grants R35GM130331 and R01NS102279) and the Fast Forward Protein Folding Disease Initiative of the University of Michigan, and both A.W.T. and Y.W. were supported by American Lung Association (Grant COVID-1019544).

## AUTHOR CONTRIBUTIONS

J.Z., A.W.T, and Y.W. designed the experiments. A.K., J.Z., and D.J. performed all the viral infection and fixation of infected cells. J.Z. prepared IF and EM samples. J.Z., W.R., J.J., L.X., M.R., and S.E. performed IF imaging and quantification. J.Z. and L.X. performed EM imaging, cell surface biotinylation assay, lentivirus preparation, and immunoblotting. S.B. and J.Z. performed immunoblotting of cell samples w/o infection. L.X. and J.Z. performed virus cell entry assay. J.Z. and D.J. performed the PEG precipitation, TCID50, and plaque assays. J.Z., A.W.T and Y.W. analyzed the data and wrote the paper, with inputs from all authors.

## DECLARATION OF INTERESTS

The authors declare no competing interests.

---

**Figure S1.**
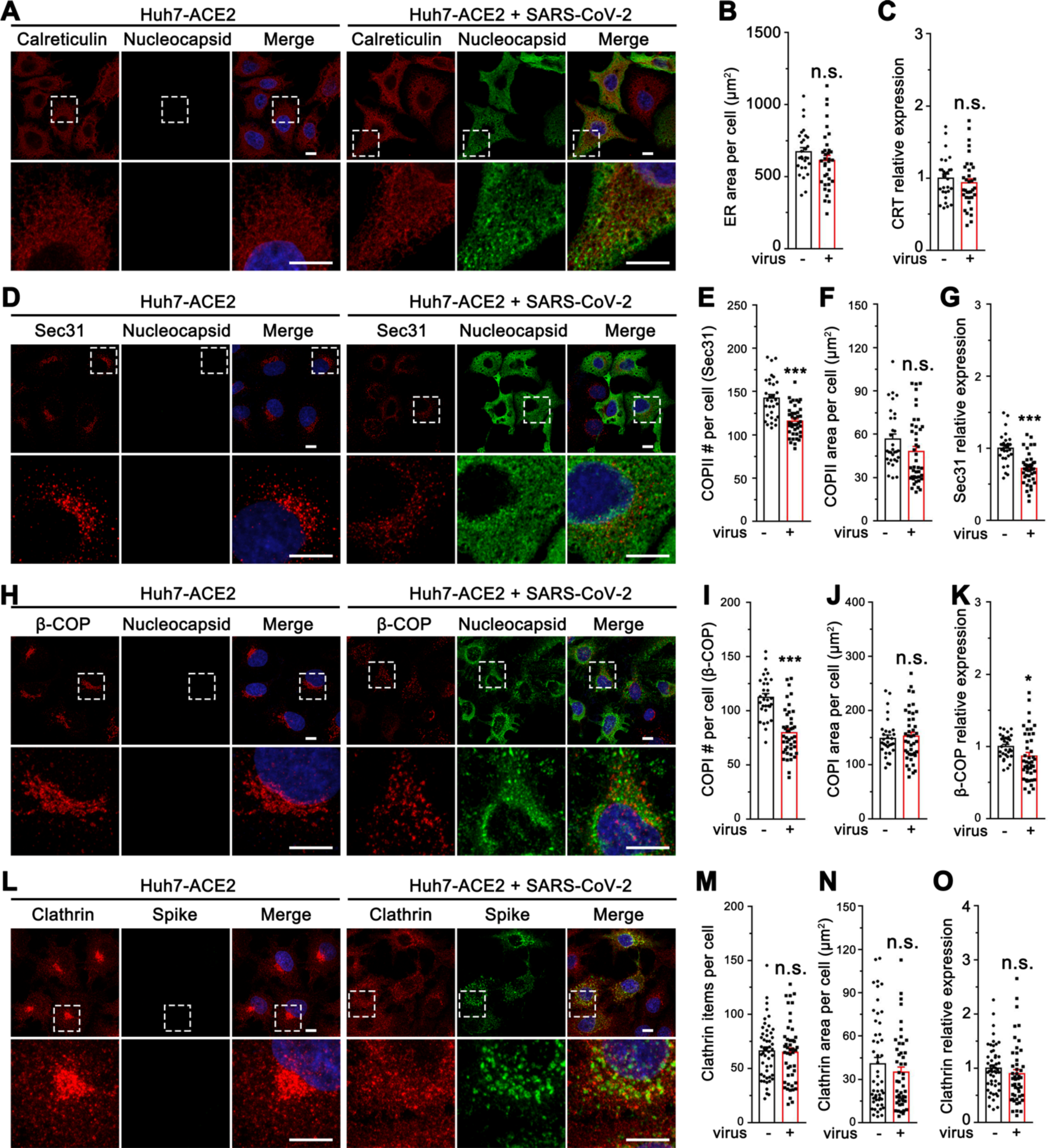
SARS-CoV-2 infection modulates the formation of COPI and COPII vesicles. (A) Representative confocal images of Huh7-ACE2 cells incubated with or without SARS-CoV-2 (MOI = 1) for 24 h and stained for calreticulin and nucleocapsid. (B-C) Quantification of calreticulin for the area (B) and relative expression (C) in A. (D) Representative confocal images of Huh7-ACE2 cells incubated with or without SARS-CoV-2 (MOI = 1) for 24 h and stained for Sec31 and nucleocapsid. (E-G) Quantification of Sec31 item number (E), area (F), and relative expression (G) in D. (H) Representative confocal images of Huh7-ACE2 cells incubated with or without SARS-CoV-2 (MOI = 1) for 24 h and stained for β-COP and spike. (I-K) Quantification of β-COP in H. (L) Representative confocal images of Huh7-ACE2 cells incubated with or without SARS-CoV-2 (MOI = 1) for 24 h and stained for clathrin and nucleocapsid. (M-O) Quantification of clathrin in L. Boxed areas in the upper panels are enlarged and shown underneath. Scale bars in all panels, 10 μm. All quantitation data are shown as mean ± SEM from three independent experiments. Statistical analyses were performed using two-tailed Student’s t-test. *, p < 0.05; ***, p < 0.001; n.s., not significant.

**Figure S2.**
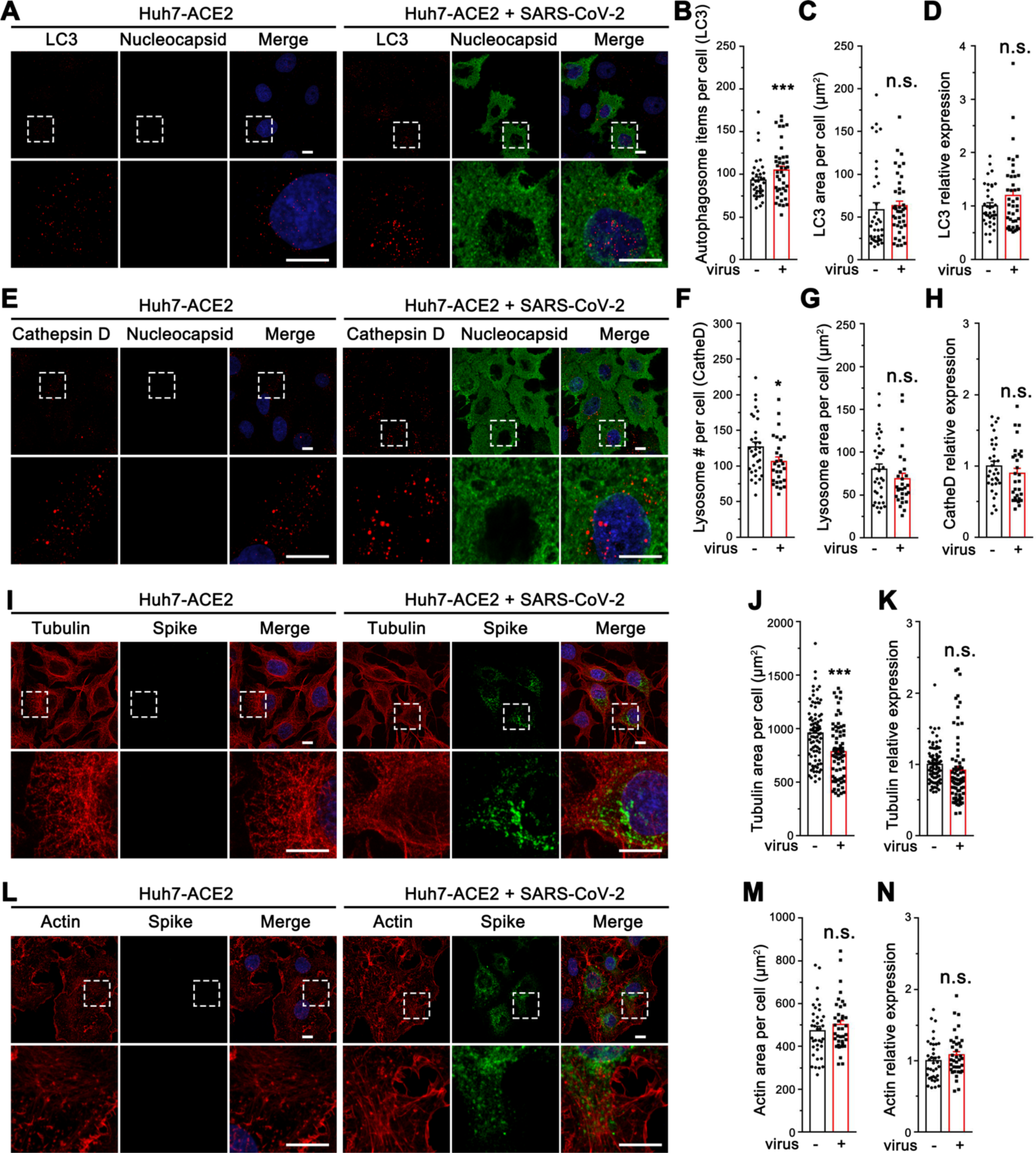
SARS-CoV-2 infection increases autophagy but exhibits minor effects on cytoskeleton. (A) Representative confocal images of Huh7-ACE2 cells incubated with or without SARS-CoV-2 (MOI = 1) for 24 h and stained for LC3 and nucleocapsid. (B-D) Quantification of LC3 item number (B), area (C), and relative expression (D) in A. (E) Representative confocal images of Huh7-ACE2 cells incubated with or without SARS-CoV-2 (MOI = 1) for 24 h and stained for cathepsin D and nucleocapsid. (F-H) Quantification of cathepsin D in E. (I) Representative confocal images of Huh7-ACE2 cells incubated with or without SARS-CoV-2 (MOI = 1) for 24 h and stained for a-Tubulin and spike. (J-K) Quantification of tubulin in I. (L) Representative confocal images of Huh7-ACE2 cells incubated with or without SARS-CoV-2 (MOI = 1) for 24 h and stained for actin (with phalloidin) and spike. (M-N) Quantification of actin in L. Boxed areas in the upper panels are enlarged and shown underneath. Scale bars in all panels, 10 μm. All quantitation data are shown as mean ± SEM from three independent experiments. Statistical analyses were performed using two-tailed Student’s t-test. *, p < 0.05; ***, p < 0.001; n.s., not significant.

**Figure S3.**
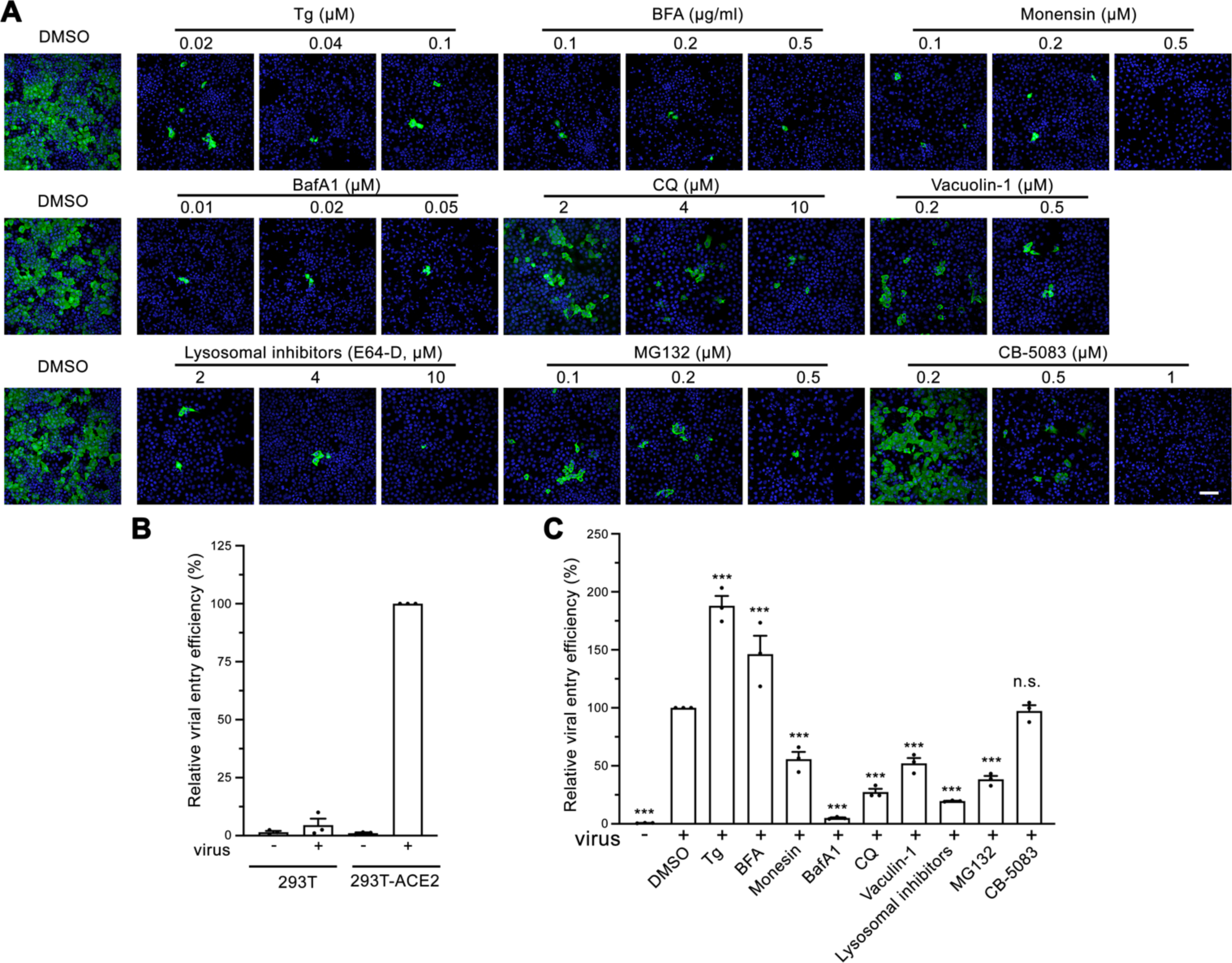
Molecules disrupting Golgi functions inhibit SARS-CoV-2 infection at low concentrations. (A) Representative confocal images of Huh7-ACE2 cells infected with SARS-CoV-2 for 24 h in the presence of indicated molecules and stained for nucleocapsid. Scale bar, 100 μm. Quantification of the viral infection rate is shown in Figure 2C. (B) Validation of cell entry assay of 293T and 293T-ACE2 cells by infection of SARS-CoV-2 Spike pseudotyped lentiviruses. (C) Cell entry assay of 293T or 293T-ACE2 cells by SARS-CoV-2 Spike pseudotyped lentivirus for 24 h in the presence of indicated molecules. Data are shown as mean ± SEM from three independent experiments. Statistical analyses are performed using One-way ANOVA. ***, p < 0.001, n.s., not significant.

**Figure S4.**
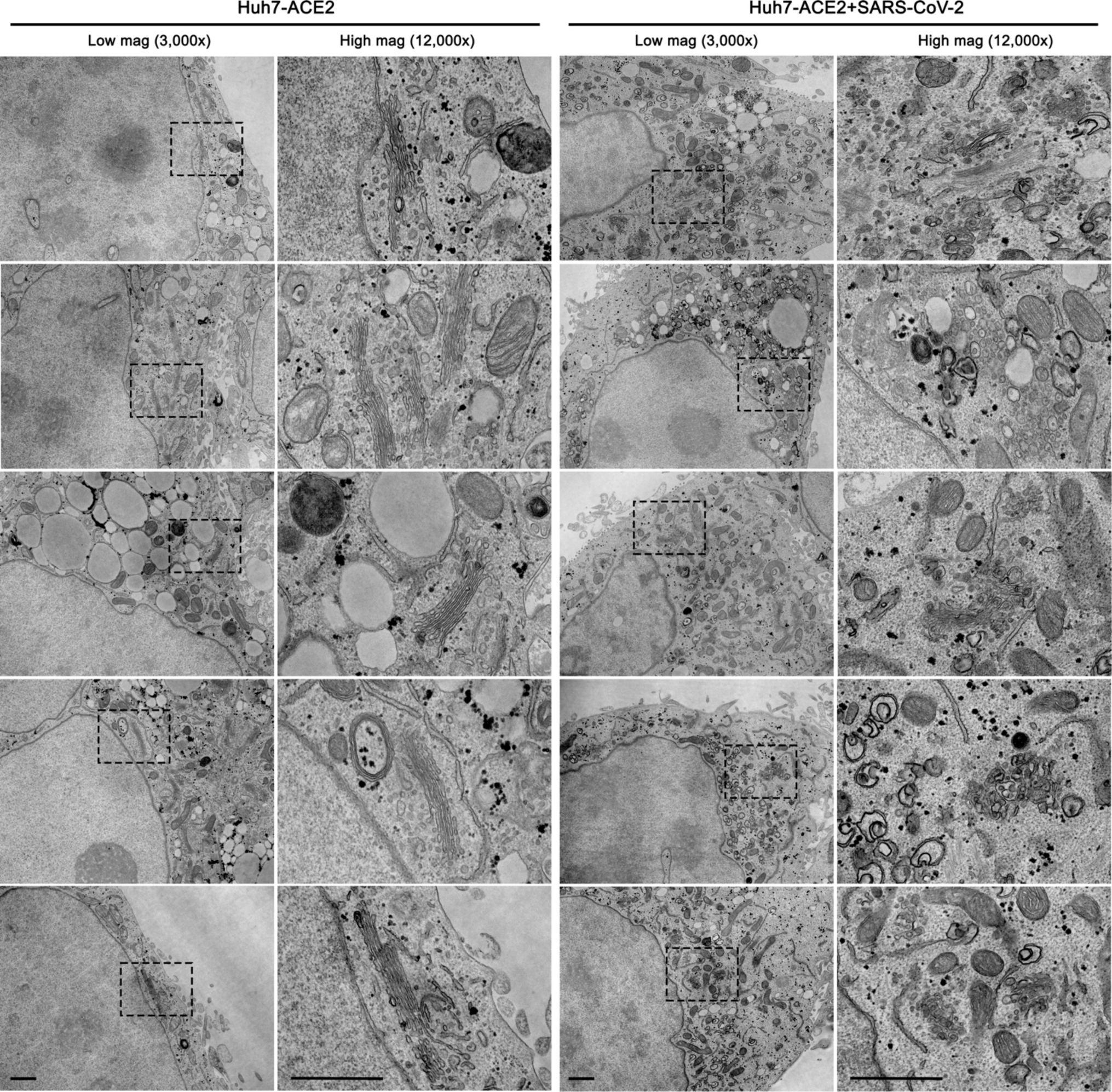
SARS-CoV-2 remodels the Golgi and viral particles accumulate in Golgi fragments. A gallery of EM images of Huh7-ACE2 cells incubated with or without SARS-CoV-2 (MOI = 1) for 24 h under two different magnifications. Boxed areas on the left images are enlarged and shown on the right. Scale bars, 500 nm.

**Figure S5.**
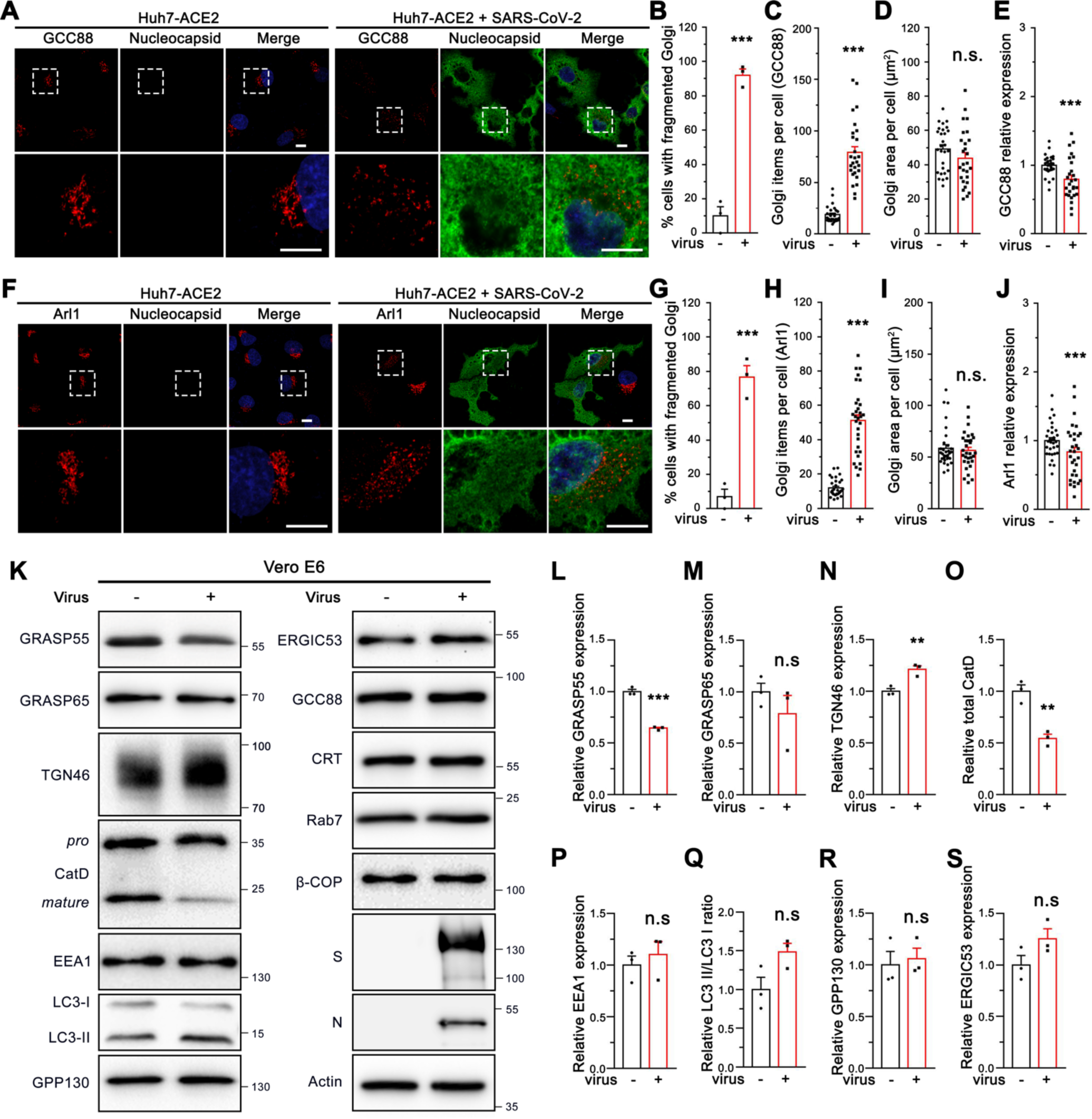
SARS-CoV-2 infection alters the Golgi structure and GRASP55 and TGN46 expression. (A) Representative confocal images of Huh7-ACE2 cells incubated with or without SARS-CoV-2 (MOI = 1) for 24 h and stained for a trans-Golgi marker GCC88 and nucleocapsid. (B-E) Quantification of A for the percentage of cells with fragmented Golgi (B), GCC88 item number (C), area (D), and relative expression level (E). (F) Representative confocal images of Huh7-ACE2 cells incubated with or without SARS-CoV-2 (MOI = 1) for 24 h and stained for a trans-Golgi marker Arl1 and nucleocapsid. Boxed areas in the upper panels of A and F are enlarged and shown underneath. Scale bars, 10 μm. (G-J) Quantification of Arl1 in F. (K) Immunoblots of indicated proteins in Vero E6 cells incubated with or without SARS-CoV-2 (MOI = 1) for 24 h. (L-S) Quantification of K for the relative level of GRASP55 (L), GRASP65 (M), TGN46 (N), cathepsin D (CatD, O), EEA1 (P), LC3-II/LC3-I ratio (Q), GPP130 (R) and ERGIC53 (S) in K. Quantitation data are shown as mean ± SEM from at least three independent experiments. Statistical analyses were performed using two-tailed Student’s t-test. **, p < 0.01; ***, p < 0.001; n.s., not significant.

**Figure S6.**
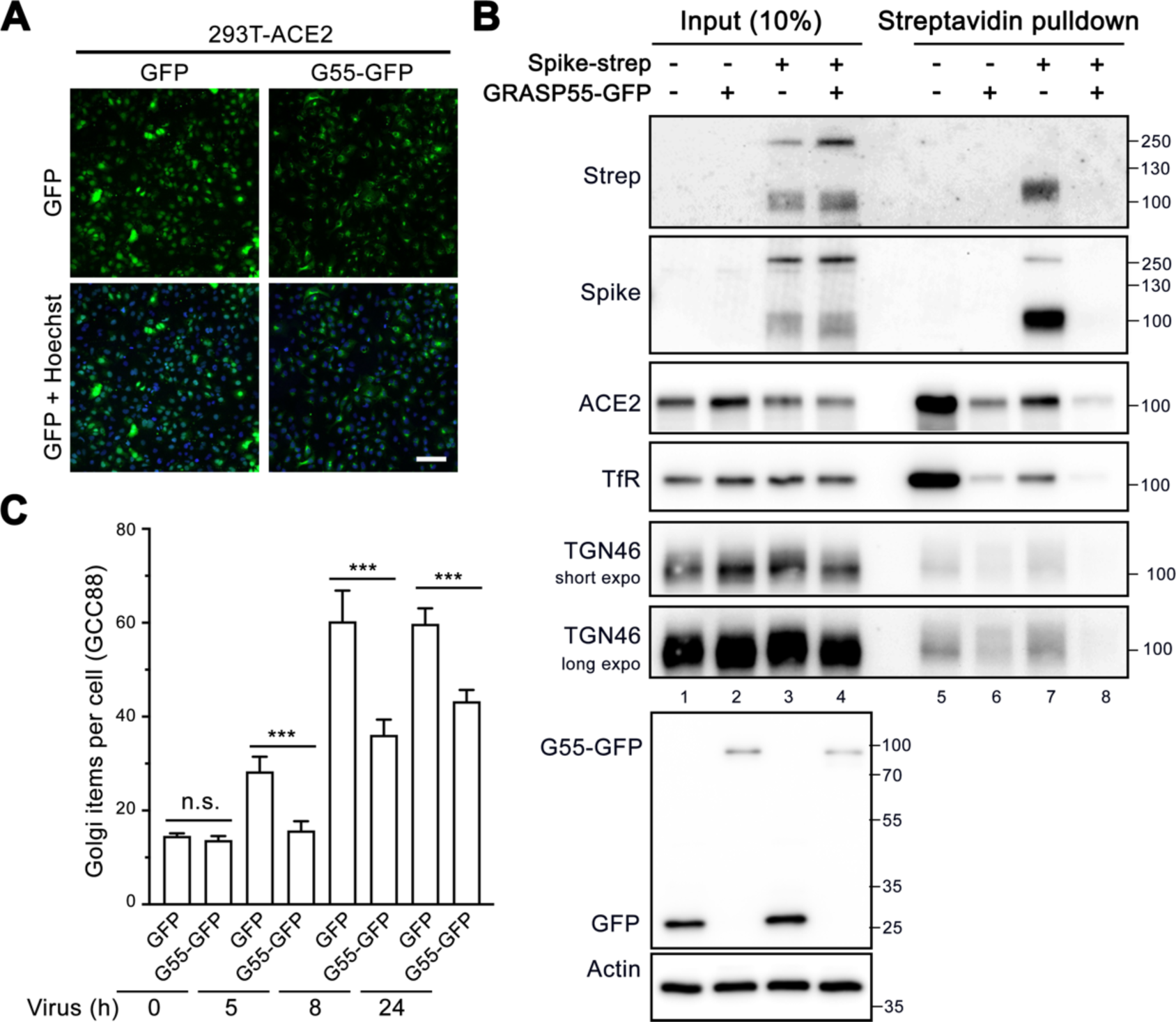
GRASP55 expression reduces spike trafficking. (A) Representative confocal images of stable 293T-ACE2 cells expressing GFP or GRASP55-GFP. Scale bar, 100 μm. (B) GRASP55 expression reduced spike at the cell surface. Huh7-ACE2 cells were transfected with GFP or GRASP55-GFP for 24 h, and then co-transfected with GFP or GRASP55-GFP together with spike-strep for 24 h. Cell surface proteins were biotinylated, pulled down by streptavidin beads, and blotted for indicated proteins. (C) Quantification of the Golgi item number of GCC88 of Huh7-ACE2 cells stably expressing GFP or GRASP55 infected by SARS-CoV-2 for 5, 8, and 24 h. Data are shown as mean ± SEM from more than 24 random images from two independent experiments. Statistical analyses were performed using two-tailed Student’s t-test. ***, p < 0.001; n.s., not significant.

**Figure S7.**
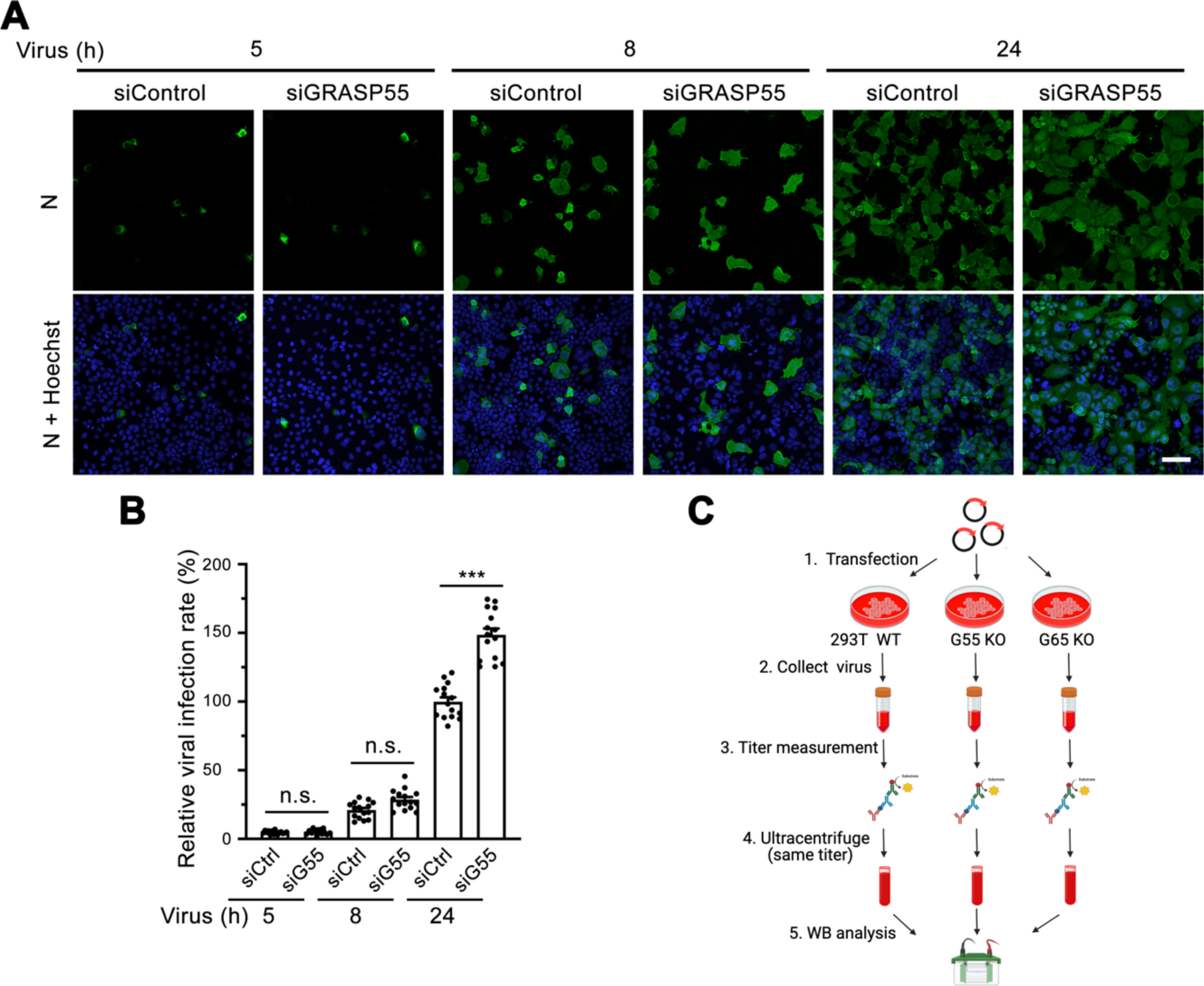
GRASP55 depletion increases viral infection at late but not early stage. (A) Representative confocal images of Huh7-ACE2 cells transfected with siControl or siGRASP55 oligos for 48 h followed by infection with SARS-CoV-2 for 5, 8, and 24 h. Scale bar, 100 μm. (B) Quantification of the viral infection rate in B. Data are shown as mean ± SEM from 30 random images from two independent experiments. Statistical analyses were performed using One-way ANOVA. ***, p < 0.001; n.s., not significant. (C) Diagram of immunoblotting analysis of same-titer SARS-CoV-2 Spike pseudotyped lentiviruses collected from wild-type, GRASP55 KO, and GRASP65 KO 293T cells in Figure 5M.

**Figure S8.**
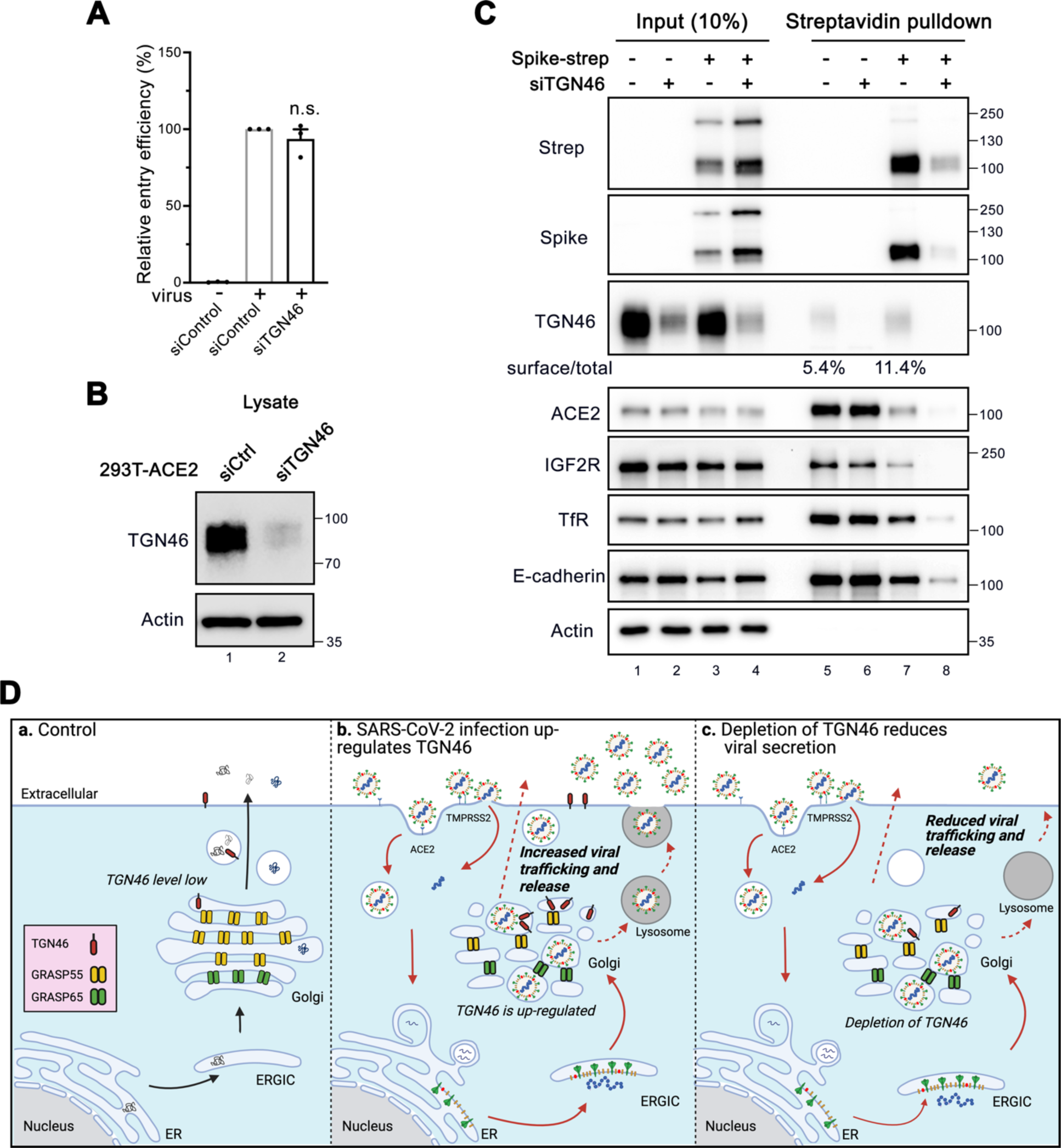
Depletion of TGN46 inhibits SARS-Co-V secretion without affecting viral entry. (A) Cell entry assay of 293T-ACE2 cells transfected with siControl or siTGN46 oligos for 48 h followed by infection with SARS-CoV-2 Spike pseudotyped lentivirus for 24 h. Statistical analyses were performed using two-tailed Student’s t-test. n.s., not significant. (B) Immunoblots of cell lysates of 293T-ACE2 cells transfected with siControl or siTGN46 oligos for 48 h followed by infection with SARS-CoV-2 Spike pseudotyped lentivirus for 24 h. (C) TGN46 depletion reduces spike protein at the cell surface. Huh7-ACE2 cells were transfected with siControl or siTGN46 oligos for 48 h followed by transfection with spike-strep for 24 h. Cell surface proteins were biotinylated, pulled down by streptavidin beads, and blotted for indicated proteins. Note that TGN46 level is enhanced at the cell surface by spike protein expression. (D) Proposed working model for a novel role of TGN46 in SARS-CoV-2 infection. In brief, under normal conditions (a) TGN46 is expressed at a relatively low level and recycles between the Golgi and plasma membrane. After SARS-CoV-2 infection (b), TGN46 is up-regulated, which accelerates viral trafficking. When TGN46 is depleted (c), the trafficking speed of all variants of SARS-CoV-2 is reduced. Thus, TGN46 may serve as a carrier for viral trafficking and release.

## Supplemental movies

**Movie S1. Intact Golgi in a control cell.**

3D reconstruction of Huh7-ACE2 cells stained for GM130. Images were taken for a total of 40 stacks and maximum intensity projection was performed.

**Movie S2. SARS-CoV-2 infection causes Golgi fragmentation.**

Huh7-ACE2 cells were infected with SARS-CoV-2 (MOI = 1) for 24 h, fixed by 4% PFA, and stained for spike and GM130. Images were taken for a total of 40 stacks and maximum intensity projection was performed. Note the colocalization of spike and GM130 in the Golgi fragments.

**Supplementary Table 1.** Antibody list used in this study.

| REAGENT or RESOURCE | SOURCE | IDENTIFIER |
| --- | --- | --- |
| Mouse monoclonal anti-SARS-CoV-2 N | ProSci | Cat# ABIN6952432 |
| Rabbit polyclonal anti-SARS-CoV-2 spike (C terminal) | ProSci | Cat# 3525 |
| Rabbit polyclonal anti-SARS-CoV-2 spike (S1) | ProSci | Cat# 9083 |
| Mouse monoclonal anti-Strep Tag | Qiagen | Cat# 34850; RRID: AB_2810987 |
| Rabbit polyclonal anti-ACE2 | Proteintech Group | Cat# 21115-1-AP; RRID: AB_10732845 |
| Rabbit polyclonal anti-Calreticulin | Proteintech Group | Cat# 27298-1-AP; RRID: AB_2880835 |
| Rabbit polyclonal anti-ERGIC53 | Proteintech Group | Cat# 13364-1-AP; RRID: AB_2135994 |
| Rabbit polyclonal anti-GPP130 | Biolegend | Cat# 923801; RRID: AB_2565442 |
| Mouse monoclonal anti-GM130 | BD Biosciences | Cat# 610823; RRID: AB_398142 |
| Rabbit polyclonal anti-GRASP65 | Gift from Joachim Seemann | N/A |
| Rabbit polyclonal anti-GRASP55 | Proteintech Group | Cat# 10598-1-AP; RRID: AB_2113473 |
| Rabbit polyclonal anti-GCC88 | Proteintech Group | Cat# 16271-1-AP; RRID: AB_2107197 |
| Rabbit polyclonal anti-Golgin-97 | Proteintech Group | Cat# 12640-1-AP; RRID: AB_2115315 |
| Rabbit polyclonal anti-GalT | Novus | Cat# NBP1-88654; RRID: AB_11018155 |
| Sheep polyclonal anti-TGN46 | Bio-Rab/AbD Serotec | Cat# AHP500; RRID: AB_324049 |
| Mouse monoclonal anti-Golgin-245 | BD Biosciences | Cat# 611280; RRID: AB_398809 |
| Rabbit monoclonal anti-Arl1 | Abcam | Cat# ab155982 |
| Rabbit polyclonal anti-β-COP | Gift from Felix Wieland | N/A |
| Rabbit polyclonal anti-Sec31 | Gift from Fred Gorelick | N/A |
| Mouse monoclonal anti-Clathrin heavy chain | Gift from Frances Brodsky | N/A |
| Mouse monoclonal anti-EEA1 | Santa Cruz Biotechnology | Cat#sc-365652; RRID: AB_10850311 |
| Rabbit polyclonal anti-Rab7 | Proteintech Group | Cat# 55469-1-AP; RRID: AB_11182831 |
| Rabbit polyclonal anti-LC3B | Cell Signaling Technology | Cat# 2775; RRID: AB_915950 |
| Rabbit polyclonal anti-Cathepsin D | Proteintech Group | Cat# 21327-1-AP; RRID: AB_10733646 |
| Mouse monoclonal anti-LAMP2 | DSHB | Cat# H4B4; RRID: AB_2134755 |
| Rabbit polyclonal anti-IGF2R | Proteintech Group | Cat# 20253-1-AP; RRID: AB_10859779 |
| Mouse monoclonal anti-TfR | Thermo Fisher | Cat# 13-6800; RRID: AB_2533029 |
| Rabbit polyclonal anti-E-cadherin | Proteintech Group | Cat# 20874-1-AP; RRID: AB_10697811 |
| Mouse monoclonal anti-alpha-Tubulin | DSHB | Cat# AA4.3; RRID: AB_579793 |
| Mouse monoclonal anti-beta-Actin | Proteintech Group | Cat# 66009-1-lg; RRID: AB_2687938 |

## Notes

### Competing Interest Statement

The authors have declared no competing interest.

### Summary of Updates

Figures 3A-E, 5, 6, 7H-N,8, Figures S3B-C, S6A, S6C, S7, S8A-B are new results.

